# Exploring Links between Brain Image-Derived Phenotypes and Accelerometer-Measured Physical Activity in the UK Biobank

**DOI:** 10.64898/2026.03.10.710798

**Authors:** Dongliang Zhang, Andrew Leroux, Ciprian M. Crainiceanu, Martin A. Lindquist

## Abstract

A broad range of neurodegenerative disorders are associated with altered functional connectivity (FC) patterns and atrophy of gray matter volume (GMV). Similarly, there are links between physical activity (PA) and a number of neurodegenerative disorders. However, studies investigating the link between brain image-derived phenotypes (IDPs) and PA remain limited. Using data from the UK Biobank, we investigated the multivariate association between two sets of brain IDPs (related to FC and GMV) and PA using canonical correlation analysis (CCA). We further quantified the importance of individual PA variables in modeling each set of IDPs using both supervised and unsupervised approaches, and assessed their predictive performance for individual brain phenotypes. Finally, we evaluated the predictive performance of brain IDPs and PA variables for diabetes, stroke, coronary heart disease (CHD), and cancer using nested logistic regression models, with their relative contributions to explained variation in disease status quantified using a coefficient of determination specifically designed for logistic regression. Our analyses identified a statistically robust but low-dimensional axis of shared variation between PA and FC (canonical correlation r = 0.50), whereas the corresponding association between PA and GMV was weaker (r = 0.19). Brain features contributing most strongly to these associations were located in motor- and attention-related networks. Across predictive models, a small set of correlated PA measures reflecting activity intensity and circadian rhythm consistently emerged as representative predictors of both FC and GMV variation. Finally, we found that PA variables demonstrated greater predictive utility than either FC or GMV alone, particularly for CHD and diabetes, as assessed by both the area under the receiver operating characteristic (ROC) curve (AUC) and the proportion of explained variation. Together, these findings indicate that objectively measured PA is strongly associated with a set of motor-related brain features and provides substantial predictive information for cardiometabolic disease risk, while cross-sectional neuroimaging measures offer more modest incremental explanatory value.

## 1 Introduction

Recent advances in neuroscience research have highlighted the intricate connections between a wide spectrum of neurological and psychiatric disorders, as well as differences in both functional and structural patterns in the brain. Specifically, extensive research has shown that disruptions in functional connectivity (FC) networks, and structural impairments such as gray matter volume (GMV) atrophy, play important roles in the progression of many brain-related disorders. These pathological changes have been implicated in Alzheimer’s disease (Delbeuck et al., 2003; Wu et al., 2021), Parkinson’s disease (Gao & Wu, 2016; Charroud & Turella, 2021), schizophrenia (Lynall et al., 2010; Vita et al., 2012), depression (Dunlop et al., 2023; Shad et al., 2012), epilepsy (Royer et al., 2022; Doucet et al., 2016), multiple sclerosis (Tahedl et al., 2018; Hulst & Geurts, 2011) and Huntington’s disease (Espinoza et al., 2018; Stoffers et al., 2010). In these conditions, disruptions in FC networks have been shown to reflect functional disintegration across critical neural systems, while GMV loss can signify irreversible structural damage. For these reasons, identifying reliable functional and structural markers is crucial for improving early detection, tracking disease trajectories, and informing therapeutic decision-making.

In parallel, a growing body of evidence supports the beneficial role of physical activity (PA) in maintaining brain health (Zhao et al., 2023; Campbell & Cullen, 2023; Zhao et al., 2024; Wanigatunga et al., 2025; Hoogen et al., 2025) and reducing disease risk (Celis-Morales et al., 2016; Barker et al., 2019; Leroux et al., 2020; Watts et al., 2023; Agarwala et al., 2024; Leroux et al., 2024). PA is widely recognized not only as a modifiable lifestyle factor that lowers the risk of several of the aforementioned neurodegenerative diseases (Islam et al., 2021; Tsukita et al., 2022), but also as a protective factor against a broader class of noncommunicable diseases, such as cardiovascular disease (Ledbetter et al., 2022), diabetes (Steeves et al., 2015), and various cancers (Schrack et al., 2017). In addition, regular PA has also been shown to positively influence brain structure, particularly in aging populations, where it is associated with preservation of GMV and enhancement of white matter integrity (Erickson et al., 2014; Meijer et al., 2020). These effects can be studied using image-derived phenotypes (IDPs) extracted from multimodal brain imaging data, making them important for understanding how PA relates to brain health.

Despite substantial evidence supporting the importance of both brain imaging measures and physical activity in understanding disease mechanisms, critical gaps remain in our understanding of their joint relationships. In particular, the complex interplay between PA and various brain IDPs has not been thoroughly examined. While individual studies have demonstrated associations between specific PA metrics and isolated brain regions or imaging modalities (Hamer et al., 2018; Brown et al., 2022; Rolls et al., 2023), comprehensive analyses capturing the multivariate relationships between PA and a wide range of functional and structural brain measures are lacking. Especially underexplored is the question of how PA, measured through high-resolution accelerometry, relates simultaneously to brain FC networks and GMV across distributed brain systems.

In addition, the predictive capacity of PA for estimating variation in these brain features remains poorly understood. The extent to which PA can predict differences in FC or GMV across individuals may provide insights into pathways through which lifestyle factors affect brain aging and disease risk. Finally, there is a pressing need for integrative frameworks that jointly consider PA and brain IDPs in the context of predicting health outcomes. To date, it has not been systematically evaluated how these variables together contribute to disease risk (e.g., diabetes, stroke, cardiovascular conditions, or neurodegeneration), in a manner that allows for quantification of their relative importance or synergistic effects.

Addressing this knowledge gap is important for advancing precision medicine approaches that incorporate lifestyle, neurobiological, and behavioral data into unified models of disease risk and progression. A more complete understanding of how PA influences brain structure and function, and how these relationships contribute to broader health outcomes, could ultimately help inform targeted interventions and health policies aimed at disease prevention and healthy living.

In this study, we address these questions by leveraging data from the UK Biobank (UKB), a large-scale, population-based prospective study consisting of 500,000 participants. In addition to extensive demographic, clinical, and lifestyle data, the UKB includes the world’s largest multi-modal imaging cohort, with the goal of collecting imaging data for approximately 100,000 participants (Miller et al., 2016; Littlejohns et al., 2020). The brain magnetic resonance imaging (MRI) protocol includes three types of structural MRI, resting-state and task-based functional MRI (fMRI), and diffusion MRI (dMRI). From this data, approximately 4350 IDPs have been generated, quantifying various aspects of brain structure and function (Alfaro-Almagro et al., 2018). This comprehensive imaging dataset, combined with rich phenotypic and health outcome data, makes the UKB an invaluable resource for studying how various lifestyle factors influence the brain.

A key additional feature of the UKB is its large-scale collection of objective PA data (Leroux et al., 2020). Using the Axivity AX3 wrist-worn triaxial accelerometer, participants’ activity patterns were estimated across 8 calendar days for over 106,000 participants. Patterns of PA estimated movement behavior using a triaxial accelerometer, enabling the derivation of minute-level movement summaries reflecting participants’ real-world behavior. Combining this data with multimodal brain imaging and health outcomes offers a unique opportunity to examine how individual differences in (in)activity are associated with brain imaging-derived phenotypes, and their potential contribution to vulnerability or resilience to neurodegenerative diseases.

Using this rich dataset, we explore three interrelated objectives to clarify the joint role of PA and brain IDPs on health and disease. First, we perform a holistic multivariate analysis using bipartial canonical correlation analysis (CCA) (Hotelling, 1936), examining the associations between 11 summary PA variables and two sets of brain IDPs. Specifically, functional connectomes derived from 55 brain components and gray matter volumes measured across 139 regions of interest (ROIs). After adjusting for potential confounders, we identified statistically significant CCA modes of population co-variation, with the primary mode linking PA to brain regions responsible for movement-related functions. These findings reveal distinct and interpretable patterns of PA, FC, and GMV variables contributing to the shared axes of variability, suggesting that movement-related brain networks are particularly associated with differences in PA.

Second, we quantify the predictive value of PA for modeling individual brain features. Using both linear regression models and random forest, we evaluate the variable importance of PA measures in predicting each individual FC and GMV measurement. Results indicate that PA is especially predictive of connectivity within motor-related networks and volumes of brain structures involved in motor function and coordination. Furthermore, a consistent subset of three PA variables emerge as key predictors across both functional and structural domains, suggesting a common behavioral signature influencing multiple brain systems.

Third, we assess the joint utility of PA and brain IDPs in predicting risk for four prevalent diseases, namely diabetes, stroke, coronary heart disease (CHD), and any type of cancer. We use nested logistic regression models consisting of PA, FC, and GMV variables, and their joint contributions, to predict the risks of the four selected diseases. Classification performance is assessed using the area under the receiver operating characteristic curve (ROC-AUC), while explanatory power is evaluated via McFadden’s pseudo *R*^2^ (McFadden, 1974). Across outcomes, PA variables consistently contribute more to disease status classification than brain IDPs, particularly for CHD and diabetes. These results underscore the potential of objective PA measurements not only as behavioral health indicators but also as powerful predictors of disease risk, independent of or complementary to neuroimaging-based biomarkers.

In summary, this study provides a comprehensive assessment of how PA relates to both functional and structural brain phenotypes, and how these variables jointly inform the risk of common diseases in a large population cohort. By integrating high-resolution behavioral, imaging, and clinical data, we advance the behavioral-physiological understanding of neurobiological pathways through which lifestyle behaviors relate to brain health and disease vulnerability.

## 2 Methods

### 2.1 The UK Biobank

The UKB is a prospective epidemiological cohort study consisting of 502,536 individuals aged 40-69 years, recruited between 2006 and 2010 from across the UK (Biobank, 2006). Approximately 88% of participants self-identify as having British ancestry. All participants consented to undergo extensive baseline assessments at one of 22 UKB assessment centers located in England, Scotland, and Wales. These baseline assessments collected a wide variety of self-reported sociodemographic, physical and mental health, lifestyle-related information, as well as numerous biological samples. A subset of participants were invited to participate in various enhancement sessions for supplemental data collection, including objective assessment of PA using wearable accelerometers, and a multi-modal imaging initiative including brain, cardiac and abdominal MRI, full body dual-energy X-ray absorptiometry (DXA), and carotid ultrasound. The different enhancement sessions assessed participants at varying temporal lags from their baseline assessment (2014-2020, imaging; 2013-2015, PA), with participants completing potentially different sessions. The UKB received ethical approval from the National Information Governance Board for Health and Social Care and the National Health Service North West Centre for Research Ethics Committee.

The UKB brain imaging protocol was performed on identical 3 Tesla Siemens Skyra scanners with a 32-channel head coil and operated daily at four imaging centers across the UK. The protocol incorporates six distinctive modalities: T1-weighted structural imaging (T1), resting-state functional MRI (rs-fMRI), task functional MRI (tfMRI), T2-weighted fluid-attenuated inversion recovery (T2-weighted FLAIR) imaging, diffusion-weighted imaging (dMRI), and susceptibility-weighted imaging (swMRI). Dedicated preprocessing pipelines were applied to each imaging modality. Here we focus on T1-weighted and rs-fMRI data, which produced the two sets of brain IDPs used in our study. A comprehensive description of the UKB neuroimaging pipelines and quality control procedures is provided in Alfaro-Almagro et al., 2018.

In the UKB PA protocol, each participant was instructed to wear an Axivity AX3 wrist-worn triaxial accelerometer over eight calendar days. The device sampled acceleration at 100 Hz with a dynamic range of *±*8g, capturing continuous temporal motion across three axes. Data preprocessing involved calibration to local gravity for error reduction, exclusion of abnormal and interrupted recordings. In addition, estimated non-wear periods were imputed using diurnal averages (average across days at the same clock time), and quality control criteria were used to determine valid data based on participant age, estimated wear time, and the sub-second acceleration recorded by the device. The subsecond-level data were summarized using the Euclidean norm minus one (ENMO) (van Hees et al., 2013) measure in 5-second epochs. Further data reduction was applied to the 5-second epoch data, estimating a variety of participant-level features associated with volume and patterns of motor activity, as well as circadian rhythmicity. Exact steps for deriving participant-level features are described in Leroux et al., 2020.

In total PA data is available for 93,370 participants. Of these 9240 participants have both PA and T1-weighted (GMV) data, and 8602 have both PA and rs-fMRI data.

### 2.2 Data Processing

#### T1 pipeline

The T1 pipeline consists of gradient distortion correction, skull stripping, and linear and non-linear registration to MNI152 standard space. Subsequent steps include tissue segmentation and structural image evaluation using normalization of atrophy-cross-sectional (SIENAX) analysis (Smith, 2012), which generates IDPs reflecting volumetric measures of various brain structure. In the present work, we focus on IDPs defined using GMV for 139 anatomical regions of interest (ROIs), defined using the Harvard-Oxford cortical and subcortical atlases, as well as the Diedrichsen cerebellar atlas. These 139 ROIs are graphically displayed in Figures S1-S3 in Section 3 of the Supplementary Materials. Following Alfaro-Almagro et al., 2021, sex, age, head size, brain positions (lateral, traverse, and longitudinal), position of scanner table, measure of head motion in T1 structural image, and intensity scaling for T1 were considered confounder variables for GMV.

#### rs-fMRI pipeline

The rs-fMRI pipeline includes grand-mean intensity normalization, high-pass temporal filtering, and artifact removal using ICA + FIX. Functional connectivity networks were derived by applying group principal component analysis to the normalized preprocessed data, yielding 1,200 spatial eigenvectors. These are then reduced via spatial ICA to 100 independent components with 490 time points. After discarding components identified as artifacts, 55 non-artifactual components were retained for downstream statistical analysis and interpreted as functional networks. A graphical depiction of the 55 networks is shown in Supplementary Figure S4. For each participant, a 55 *×* 55 netmat (network matrices) (Alfaro-Almagro et al., 2018), representing partial correlation between nodes, is obtained and converted into a vector representing the 1485 nodewise connections with diagonal elements excluded. An element-wise Fisher transformation is applied to each element. Following Alfaro-Almagro et al., 2021, the variables sex, age, head size, brain positions (lateral, traverse and longitudinal), position of scanner table, mean rs-fMRI head motion averaged across space and time, median absolute head motion from rs-fMRI and intensity scaling for rs-fMRI were considered confounders.

#### PA pipeline

Following the approach proposed by Leroux et al., 2020, for processing PA data in UKB, the initial five-second epoch acceleration data is averaged at minute-level for computational purposes. To capture the volume, fragmentation and circadian rhythm of PA, the following 11 variables were extracted from the summarized minute-level acceleration data: (1) total log acceleration (TLAC); (2) total minutes of light-intensity physical activity (LIPA); (3) total minutes of moderate/vigorous physical activity (MVPA); (4) sedentary bout (SBout); (5) active bout (ABout); (6) sedentary-to-active transition probability (SATP); (7) active-to-sedentary-transition probability (ASTP); (8) daytime activity ratio estimate (DARE); (9) average logarithm of acceleration during the ten most active hours of the day (M10); (10) average logarithm of acceleration during the five least active hours of the day (L5); and (11) relative amplitude (RA) defined by RA=(M10-L5)/(M10+L5). Here, the cut-off point separating LIPA from MVPA was 193 milli-g per minute, which was derived through performing quantile-matching of the distribution of MVPA time in the UKB to that recorded in the NHANES 2003-2006 dataset (Meng et al., 2023). The variables sex, age, BMI, smoking status, alcohol drinking frequency, long-standing illness status and disease history of diabetes, stroke, coronary heart disease (CHD) and cancer were considered confounder variables.

#### Disease Status

The variable **y**_(disease)_ denotes the 0-1 binary response vector representing the status of a given disease from diabetes, stroke, CHD and cancer, where 1 indicates having the disease and 0 indicates otherwise. The disease status is assigned based on whether diagnosis or specific symptoms of the disease were reported prior to the start time of recording physical activity. Specifically, a participant is considered having (1) diabetes if any type of diabetes mellitus (insulin-dependent, non-insulin-dependent, malnutrition-related or others) was reported; (2) stroke if retinal vascular occlusions, cerebral infarction, stroke (not specified as haemorrhage or infarction), intracerebral haemorrhage, subarachnoid haemorrhage and transient cerebral ischaemic attacks and related syndromes were reported; (3) CHD if chronic ischaemic heart disease was reported; (4) cancer if any type of cancer was reported.

### 2.3 Multivariate Association Analysis

Given the high dimensionality and correlations among both brain IDPs and PA variables, CCA is used here as a descriptive multivariate tool to identify low-dimensional patterns of shared covariance, rather than as a complete decomposition of dependence between the two data blocks. In particular, we emphasize interpretation of the primary mode of co-variation, which captures the dominant axis of shared structure and is the most statistically and conceptually stable in this setting.

To be specific, CCA was used to assess the multivariate symmetric association between (*i*) FC and PA variables, and (*ii*) GMV and PA variables. Ignoring the effects of confounder variables in CCA can lead to unreliable conclusions (Winkler et al., 2020). Therefore, principled selection of confounder variables for FC, GMV, and PA variables was performed, and their effects were removed before performing CCA. We performed a pairwise complete case analysis, restricting the analytic sample to participants with complete data on all the variables considered. Since imaging and PA data were collected as separate UKB enhancements, this resulted in a total of 8,580 participants for the study of CCA between FC and PA, and 9,208 participants for the study of CCA between GMV and PA. Below, we summarize the key components of the CCA methodology and the quantities reported. A more detailed illustration of the method and associated analyses is provided in Section 7 of the Supplementary Materials.

CCA identifies linear combinations of two multivariate datasets that maximize their correlation (Hotelling, 1936). While CCA yields a sequence of canonical modes, later modes are often unstable and difficult to interpret in high-dimensional, correlated settings, and typically explain minimal additional shared variance. Consequently, our inferential and interpretive focus is restricted to the primary canonical mode.

In the presence of confounding variables for each dataset, CCA is commonly performed on residualized data obtained after regressing out confounder effects, a procedure often referred to as bipartial CCA. Statistical significance of CCA modes was assessed using both asymptotic likelihood-based tests and permutation testing. However, when bipartial CCA is used, permutation tests may violate exchangeability assumptions due to residualization, potentially compromising valid inference beyond the first mode (Winkler et al., 2020). Although projection-based approaches have been proposed to restore exchangeability, numerical instability associated with singular confounder matrices precluded their reliable implementation in this dataset. To avoid over-interpretation under uncertain inferential conditions, we therefore adopt a conservative strategy and restrict interpretation to the primary CCA mode, which was consistently significant across testing approaches.

In addition to canonical correlations, we report the proportion of variance explained by each canonical variate within each data block, as well as correlations between individual PA variables and the primary canonical variate. Because CCA maximizes correlation rather than explained variance, a high canonical correlation does not necessarily imply that a large proportion of variance is shared between datasets, particularly in high-dimensional settings such as functional connectivity. Accordingly, variance-explained metrics are reported alongside canonical correlations to contextualize the scope of the identified associations.

### 2.4 Variable Importance Assessment

To determine the importance of each individual PA variable in modeling the two sets of brain IDPs (FC and GMV), the response vector is assumed to be either the 1485 nodewise FC measures between the 55 components or the GMVs from the 139 ROIs, while the design matrices contain both confounders for each data type and PA variables. Analogous to the data processing procedure for CCA, missing observations are discarded, and participants are included in the analytic sample only if they have complete data on all variables (PA, FC, GMV, confounders).

Let the matrix of response vectors be **Y** *∈* ℝ^*n×q*^ . Suppose that the design matrix 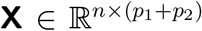 is defined via columnwise concatenation of the two matrices as **X** = [**G H**], where 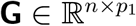 denotes the matrix of confounders and 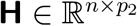 the matrix of PA variables. We focus on evaluating the statistical significance of individual predictor of **H** in modeling each of the *q* columns of **Y** and the associated predictive performance.

#### Test of Significance of Linear Modeling

Denote the columns of **Y** as 𝕐_*l*_ *∈* ℝ^*n*^, *l* = 1, …, *q*. The relation between **Y** and **X** is estimated via *q* separate linear regression models treating 𝕐_*l*_ as the response vector and **X** as the design matrix, *l* = 1, …, *q*. The *p*-values for testing the significance of the *q* linear regression models are adjusted for multiplicity using the Benjamini-Hochberg method (Benjamini & Hochberg, 1995). This step identifies an index set 𝒥 *⊆ {*1, …, *q}* such that 𝕐_*l*_ for *l* ∈ 𝒥 are significantly related to **H**, after controlling for confounders.

#### Predictive Performance

For each *l ∈* 𝒥, 10-fold cross-validation was used to obtain the predicted response vector 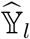. Predictive performance is evaluated by computing the Pearson’s correlation between the observed and predicted responses. This process is repeated 100 times, giving an average Pearson’s correlation between 𝕐_*l*_ and 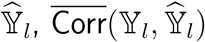, for *l ∈*𝒥 .

#### Significance of PA Variables

The variable importance of each PA variable is quantified using three methods. First, for each of the |*𝒥* | linear regression models, the significance of individual PA variables in **H** is determined using a *t*-test. Second, for *l ∈𝒥*, key PA variables are selected using stepwise BIC (both forward and backward). Third, for each *l* = 1, …, *q*, the PA variable importance is determined using random forest, which is a nonparametric, tree-based approach that estimates variable importance through evaluating each predictor’s contribution to improving prediction accuracy across an ensemble of bootstrapped tree structures. This method ultimately assigns a quantitative score to each PA variable related to its importance, providing a robust and rigorous framework for variable importance assessment by averaging across numerous decorrelated decision trees, thus leading to importance measures that are stable and capable of capturing complex nonlinear and interaction effects.

### 2.5 Disease Risk Prediction

We assessed the joint utility of PA and brain IDPs in predicting risk for four prevalent diseases, namely diabetes, stroke, coronary heart disease, and cancer. Specifically, we considered a series of nested logistic regression models in which the predictors consist of PA variables, each individual brain IDP set, and their joint specification, to model the binary disease status for each of the four selected conditions.

#### Risk Prediction

Let ***y***_disease_ *∈* ℝ^*n*^ denote the 0-1 binary disease status where 1 indicates positive disease status. Next, let the matrices of confounders, PA variables and brain IDP be respectively denoted by 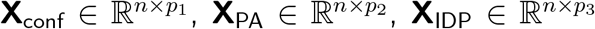. After adjusting for confounders, prediction of ***y***_disease_ is performed by fitting three nested logistic regression models to the data. These three models and the respective design matrices are given as follows:

- *M*_1_ (full model): *X*(*M*_1_) = [**X**_conf_ **X**_PA_ **X**_IDP_],
- *M*_2_ (submodel containing PA variables only): *X*(*M*_2_) = [**X**_conf_ **X**_PA_], and
- *M*_3_ (submodel containing brain IDP only): *X*(*M*_3_) = [**X**_conf_ **X**_IDP_].

We used 10-fold cross-validation (CV) to investigate and compare the binary classification performance of the three nested models, for each disease. We computed various predictive performance metrics, including probability of having the disease, the associated false positive and true positive rates, and the area under the ROC curve (AUC). While cross-validation addresses issues in predictive modeling related to potential model overfit and generalizability, variability in estimated 10-fold cross-validation arises from the potential different test/train splits of the data. Here, we repeated 10-fold cross-validation 100 times using random splits of the data, averaging over the replications to obtain final estimates of cross-validated model performance. Specifically, separately for each replication, we estimated a single ROC curve using predicted for risk for each of the 10 folds. Next, these 100 ROC curves are then averaged across replications. Finally, AUC is calculated using numerical integration of this mean ROC curve. We note that this procedure is equivalent to estimating AUC within each of the replicates and then averaging those quantities (Chen & Samuelson, 2014).

#### Proportion of Explained Variation

Based on the population-level summaries shown in Supplementary Table S4, the prevalence of diabetes, stroke, CHD and cancer for the UKB participants included in this particular study are 4.5%, 2.3%, 3.3% and 12.5%, respectively, leading to imbalanced datasets. Moreover, the disease prevalence for diabetes (Zhou et al., 2024), stroke (Imoisili et al., 2024), CHD (Lee et al., 2022), and cancer (Bizuayehu et al., 2024) is relatively low in the general population. This imbalance, along with discrepancies in the prevalence within the conditions investigated in our study, has important implications for the interpretation of AUC. In fact, even in the case of balanced datasets, ROC-based classification accuracy is seen as a discontinuous improper scoring rule, which may not serve as a reliable performance benchmark (Harrell, 2025). Therefore, in addition to computing the AUC as an initial diagnostic, we consider a likelihood-based assessment metric that bypasses the use of thresholds for predicted probabilities, providing an alternative evaluation measure of prediction performance. McFadden’s pseudo *R*^2^ (McFadden, 1974) represents the proportion of variation in the categorical data explained by a model, a commonly used benchmark in public health and social science research (Matthay et al., 2021) which is straightforward to interpret, comparatively stable across sample sizes, and well-suited for evaluating logistic and other nonlinear models. To be specific, when modeling the 0-1 binary status of a disease, this metric is computed as 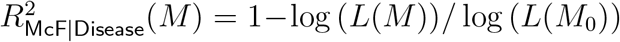, where log (*L*(*M*)) denotes the logarithm of likelihood obtained based on the fitting model *M* and log (*L*(*M*_0_)) is the logarithm of likelihood obtained based on the null model consisting of confounders only. This metric is repeatedly calculated based on the 100 testing datasets used in our repeated cross-validation procedure. The average 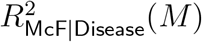 across test/train splits is reported.

## 3 Results

### Data processing for CCA

The flow charts of preparing and processing data involved in using CCA to determine the association between PA and each of the two IDPs (FC and GMV) are shown in Supplementary Figures S5 and S6, respectively. Additionally, population characteristics of the two subsets UKB participants who contributed to the two CCAs (PA/FC, PA/GMV) are summarized in Tables S1-S3 in Section 1 of the Supplementary Materials. Among the 8580 and 9208 participants respectively included in the CCA relating FC to PA and GMV to PA, we observed an overlap of 8580 participants who were included in both sets of analyses. The population-level functional connectivity (prior to Fisher transformation) is given in Figure S11(A) from Section 5 of the Supplementary Materials. Population characteristics of various confounders and PA variables used in the bipartial CCA relating PA variables to both brain IDPs are similar, despite differing sample sizes.

### Data processing for variable importance assessment

The flow charts of processing data included in determining the variable importance of individual PA variables in modeling FC and GMV are shown in Supplementary Figures S7 and S8, respectively. The population statistics for the subset of the UKB participants involved in our study are summarized in Tables S4 and S5 from Section 2 of the Supplementary Materials. The brain connectivity pattern (prior to Fisher transformation) is shown in Figure S11(B) from Section 5 of the Supplementary Materials. Analogous to the association analysis, almost identical patterns in the summary statistics are detected with respect to both IDPs. Moreover, the FC pattern in Figure S11(B) closely resembles its counterpart in Figure S11(A).

### Data processing for disease risk prediction

The data processing flow charts for predicting the risks of diabetes, stroke, CHD and cancer through three nested logistic regression models consisting of PA variables and brain IDPs are shown in Figures S9 and S10 from Section 4 of the Supplementary Materials for FC and GMV, respectively. Since the datasets involved in the risk prediction study are extracted from those used to assess variable importance, we refer to Tables S7 and S8 from Section 2 of the Supplementary Materials and Figure S11(B) for population-level summaries.

### 3.1 Multivariate Association: Functional Connectivity and Physical Activity

As described in Section 2.3, bipartial CCA was applied to the residualized physical activity 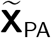 and functional connectivity 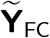, which are obtained after regressing out confounder effects from the PA variables **X**_PA_ and the FC IDP **Y**_FC_ respectively.

#### Canonical Correlations and Significance Test

Plots of the sample canonical correlation, representing the sequentially maximized correlations between linear combinations of the PA variables and the FC IDP and denoted by 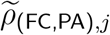 for *j* = 1, …, 11, together with the corresponding *p*-values for testing the significance of CCA modes, are shown in Figure 1 (A) and (B). Using the approximation test, the first three CCA modes of co-variation are statistically significant (*p*-values: 0, 4.14 *×* 10^*−*6^, 3.34 *×* 10^*−*3^), while all eleven are significant using permutation tests (all *p*-values approximately zero). However, due to the limitations of both tests, we focus on the primary (first) CCA mode (see Section 2.3 for discussion), whose canonical correlation 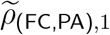 is 0.5 and *p*-values of both tests approximately equal zero.

**Figure 1.**
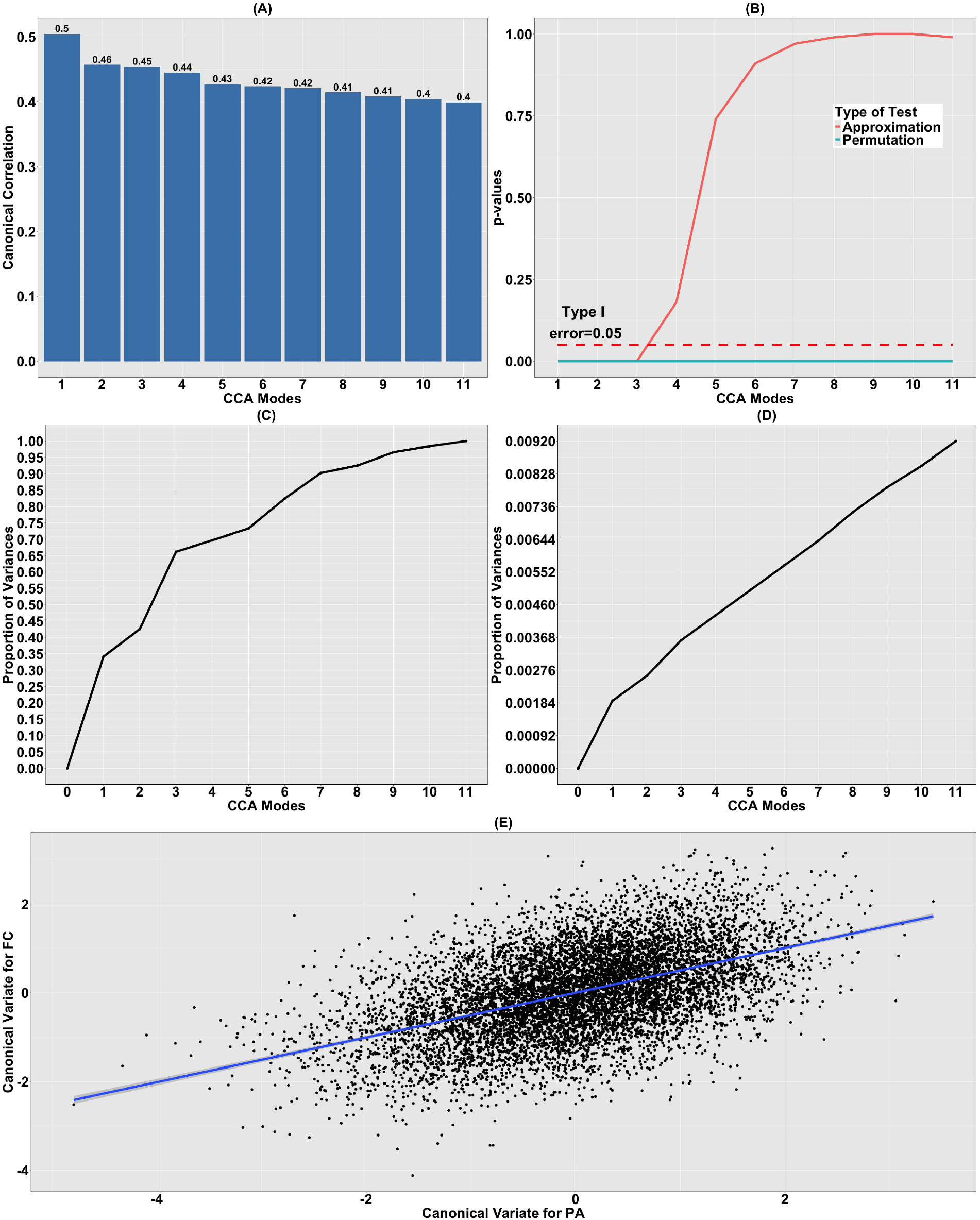
CCA relating FC to PA variables. (A) Canonical correlations of CCA relating the residualized PA variables 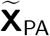 to the residualized FC variables 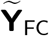; (B) corresponding p-values for testing the significance of the CCA modes using both approximation and permutation tests; (C) the cumulative proportion of variation in 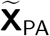 explained by 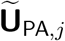, *j* = 1, …, 11; (D) the cumulative proportion of variation in 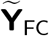 explained by 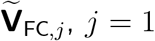, *j* = 1, …, 11; (E) the canonical variates 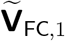 plotted against 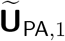. The solid blue line shows the results of a linear regression model.

#### Contribution of the CCA Modes

The contribution of each CCA mode to the variation in the datasets is visualized in Figure 1 (C) and (D), where considerable discrepancy in the contributions of sample canonical variates 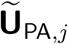 and 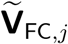 are recognized for *j* = 1, …, 11. Here, 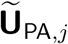 and 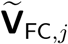 denote the canonical variates corresponding to the *j*^th^ CCA mode, representing the linear combinations of the PA variables and the FC IDPs, respectively, that are estimated to achieve the maximum sample correlation. While 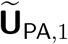 explains 34.1% of the sample variation in 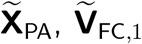 only accounts for 0.19% of the variability in 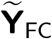. Moreover, the initial three canonical variates 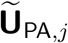 explain more than 65% of the variation. In contrast, the contribution of each canonical variate for functional connectome 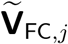 is less than 1% and subtle successive increment from 0.19% to 0.92% is seen as additional variates are included. Note that while the primary canonical variate explains a substantial proportion of variance in the PA variables, it accounts for less than 1% of total variance in the functional connectome. This indicates that the observed associations reflect a narrow subspace of brain variation aligned with dominant PA dimensions, rather than a widespread reorganization of brain connectivity or structure.

#### Canonical Variates

The sample canonical variates corresponding to the primary CCA mode, 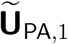 and 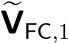, are plotted against each other in Figure 1 (E). A strong association is observed between the two data types, substantiating the significance of the primary CCA mode. This is indeed verified via fitting a linear regression model to the canonical variates, where the associated adjusted coefficient of determination of 0.25 highlights this association.

#### Canonical Coefficients for FC

To assess the contribution of each of the 1485 residualized FC measures to the primary CCA mode 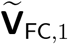, the 30 largest elements in absolute magnitude of the corresponding canonical coefficient 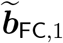 are shown in Figure 2 (A) and (B). Here, 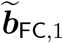 represents the estimated coefficients in the linear combination of the FC IDP that maximizes the sample canonical correlation for the primary CCA mode. Based on the figure, 14 out of the 30 edges feature positive connections, where 13 of them (92.6%) exhibit linkages within and in-between the control and attention, cingulo-opercular and sensorimotor networks. Indeed, the strongest positive connection (Z-statistic obtained via Fisher’s transformation: 1.22) bridges two components located in the control and attention and sensorimotor networks, emphasizing the high degree of relevance of these two networks to the association with PA. Considering the 16 edges with negative connections, 8 (50%) and 3 (18.8%) are between the control and attention and sensorimotor, and control and attention and cingulo-opercular networks, respectively. The edge with the strongest negative connection (Z-statistic obtained via Fisher’s transformation: -1.33) links the visual and sensorimotor networks. As such, we can see that inter- and intra-connections between control and attention and sensorimotor networks play key roles contributing to the canonical variate of the primary CCA mode 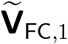 when analyzing the link between functional connectomes and physical activities, which is reasonably anticipated since these networks control and influence movement.

**Figure 2.**
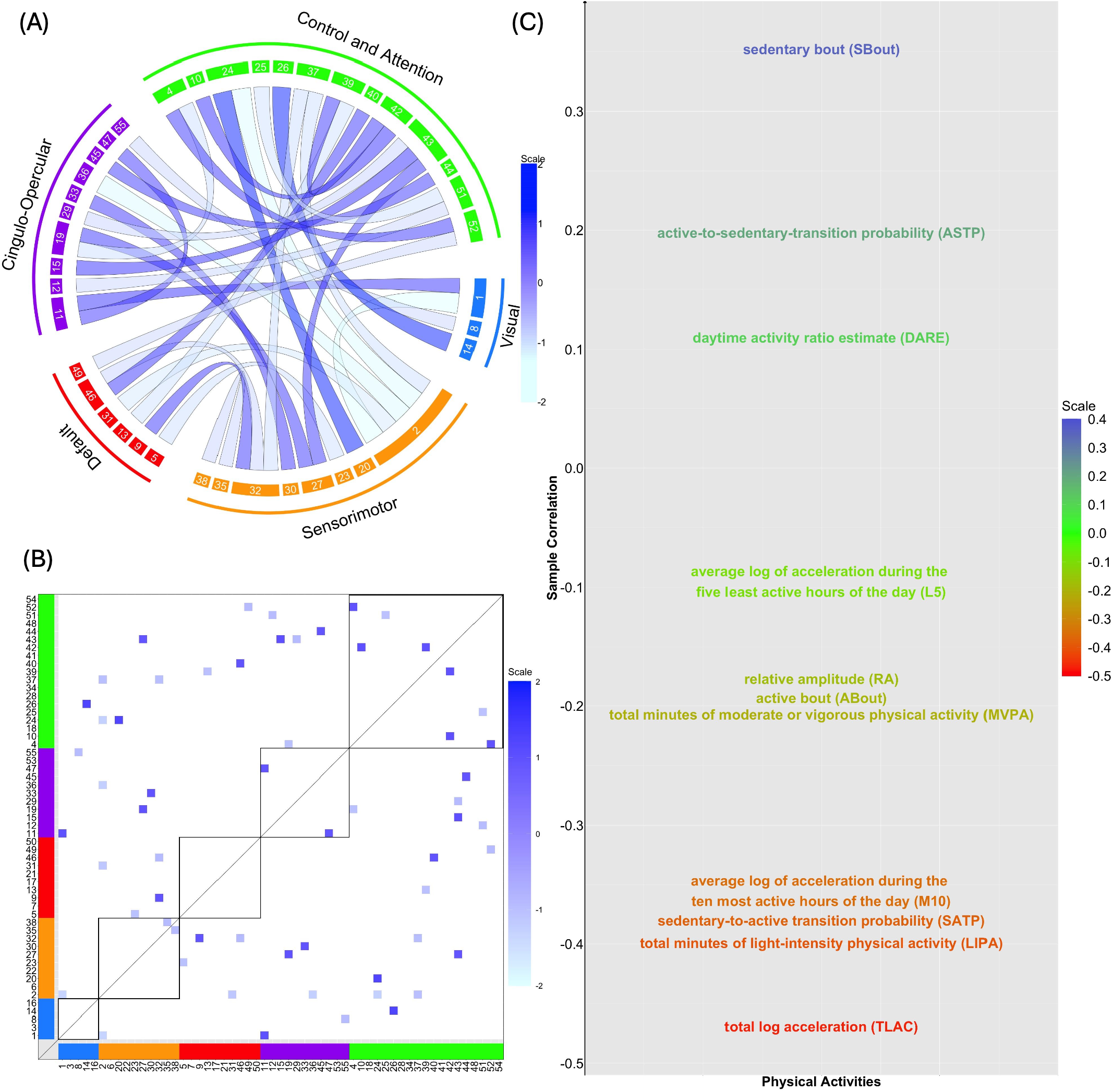
CCA relating FC to PA variables. (A) Connectogram corresponding to the 30 largest elements in absolute scale of 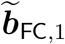; (B) same results depicted in a functional connectivity matrix; (C) sample correlations between each residualized PA variable (i.e. each column of 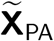) and the canonical variate for the functional connectomes 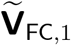 of the primary CCA mode.

#### Connection between PA variables and the Canonical Variate 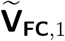

To examine the connection between each (residualized) PA variables and the identified primary CCA mode for the functional connectomes 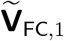, we obtain the sample correlation between each of the 11 columns of 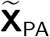 and 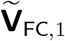, shown in Figure 2 (C). From this illustration, three of the 11 PA variables exhibit positive correlations (SBout, ASTP, and DARE) while the remaining eight exhibit negative correlations. In particular, the variable SBout, measuring an “inactive” component, is positively associated (sample correlation: 0.35) with 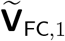. In contrast, four PA variables (TLAC, LIPA, SATP, and M10) corresponding to the volume, fragmentation and circadian rhythm of PA (Leroux et al., 2020), all considered to reflect higher activity levels, are negatively correlated (sample correlation: -0.47, -0.40, -0.38 and -0.36) with 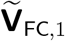. This pattern likely reflects multicollinearity among the PA variables, as sedentary measures are negatively correlated with activity-based metrics.

### 3.2 Multivariate Association: Gray Matter Volume and Physical Activity

As described in Section 2.3, bipartial CCA was applied to residualized physical activity 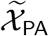 and gray matter volume 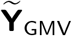, which are obtained after regressing out confounder effects from the PA variables *X*_PA_ and the GMV IDP **Y**_GMV_ respectively.

#### Canonical Correlations and Significance Test

Plots of the canonical correlations, representing the sequentially maximized correlations between linear combinations of the PA variables and the GMV IDP and denoted by 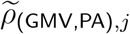 for *j* = 1, …, 11, together with the corresponding *p*-values for testing the significance of the 11 CCA modes are given in Figure 3 (A) and (B). Using the approximation, the first four modes of co-variation are statistically significant (*p*-values: 2.02 *×* 10^*−*13^, 6.11 *×* 10^*−*6^, 1.75 *×* 10^*−*3^ and 2.70 *×* 10^*−*2^), while using the permutation procedure the first ten modes are significant. Considering Winkler et al., 2020, our analysis is focused on the primary CCA mode, where 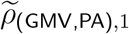 is 0.19.

**Figure 3.**
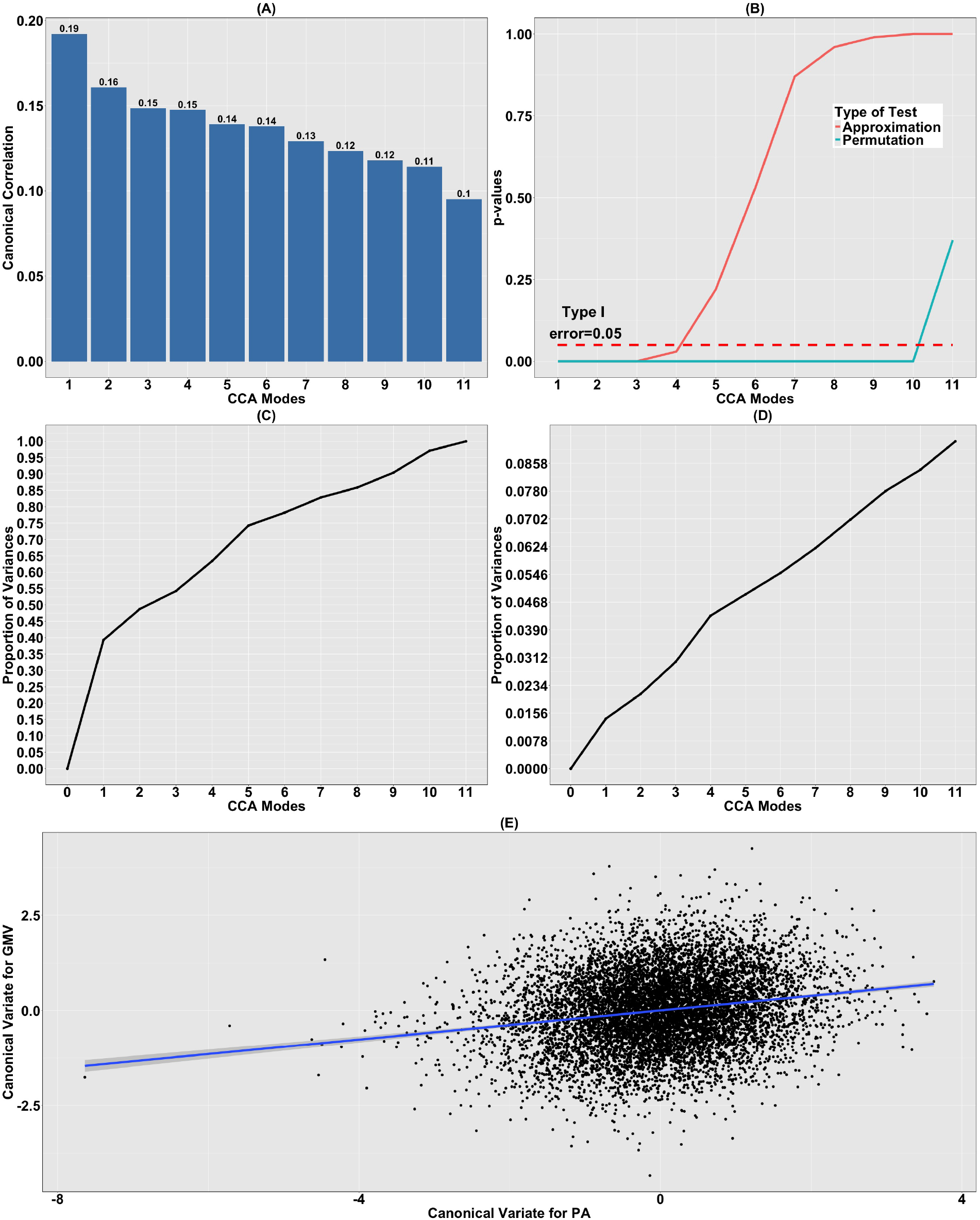
CCA relating GMV to PA variables. (A) Canonical correlations of CCA relating the residualized PA variables 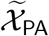 to the residualized GMV 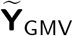; (B) corresponding p-values for testing the significance of the CCA modes using both approximation and permutation tests; (C) the cumulative proportion of variation in 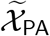 explained by 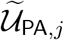, *j* = 1, …, 11; (D) the cumulative proportion of variation in 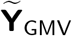 explained by 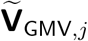, *j* = 1, …, 11; (E) plot of the canonical variates 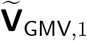 against 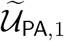. The solid blue line shows the results of a linear regression model.

#### Contribution of the CCA Modes

The contribution of each CCA mode to the variation in the datasets is visualized in Figure 3 (C) and (D). Similar to Figure 1 (C) and (D), discrepancy in the contributions of the sample canonical variates 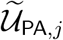 and 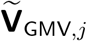 is detected, *j* = 1, …, 11. Here, 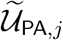 and 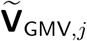 denote the canonical variates corresponding to the *j*^th^ CCA mode, representing the linear combinations of the PA variables and the GMV IDP, respectively, that are estimated to achieve the maximum sample correlation. To be more specific, 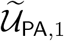 explains approximately 40% of the variation in the PA data, while 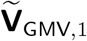 only accounts for about 1.40% for the GMV data. Moreover, from the second CCA mode onward, the successive contribution of each 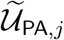 to the variation explanation slowly declines from 9.44% to 2.88%. In contrast, the variation in 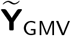 due to each 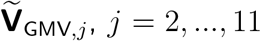, *j* = 2, …, 11 is stably maintained at about 0.7%; see Figure 3 (C) and (D). The primary canonical variate accounts for 1.4% of total variance in GMV, thus similarly to FC pointing to a narrow subspace of brain variation aligned with dominant PA dimensions.

#### Canonical Variates

The sample canonical variates corresponding to the primary CCA mode, 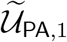 and 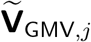, are plotted against each other in Figure 3 (E). A weaker association is detected between these two canonical variates than those observed when 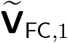 is plotted against 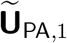 (Figure 1 (E)). This indicates a relatively weaker association between residualized PA variables and GMVs, which is further supported by the small adjusted coefficient of determination (0.04) when a linear regression model is fit to the canonical variates.

#### Canonical Coefficients for GMV

To ascertain the contribution of the GMVs for each of the 139 ROIs to the canonical variate 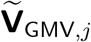, the 30 largest elements of 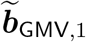 in absolute magnitude are shown in Figure 4(B). Here, 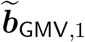 represents the estimated coefficients in the linear combination of the GMV IDP that maximizes the sample canonical correlation for the primary CCA mode. One salient feature is that all 30 coefficient components are small, ranging between -0.062 and 0.004. Specifically, these two values are attained by the GMVs corresponding to brain regions from the cerebellar vermis, a brain structure widely known to be associated with body posture, coordination of movement and locomotion. This phenomenon further highlights the significance of this region in connection with PA.

**Figure 4.**
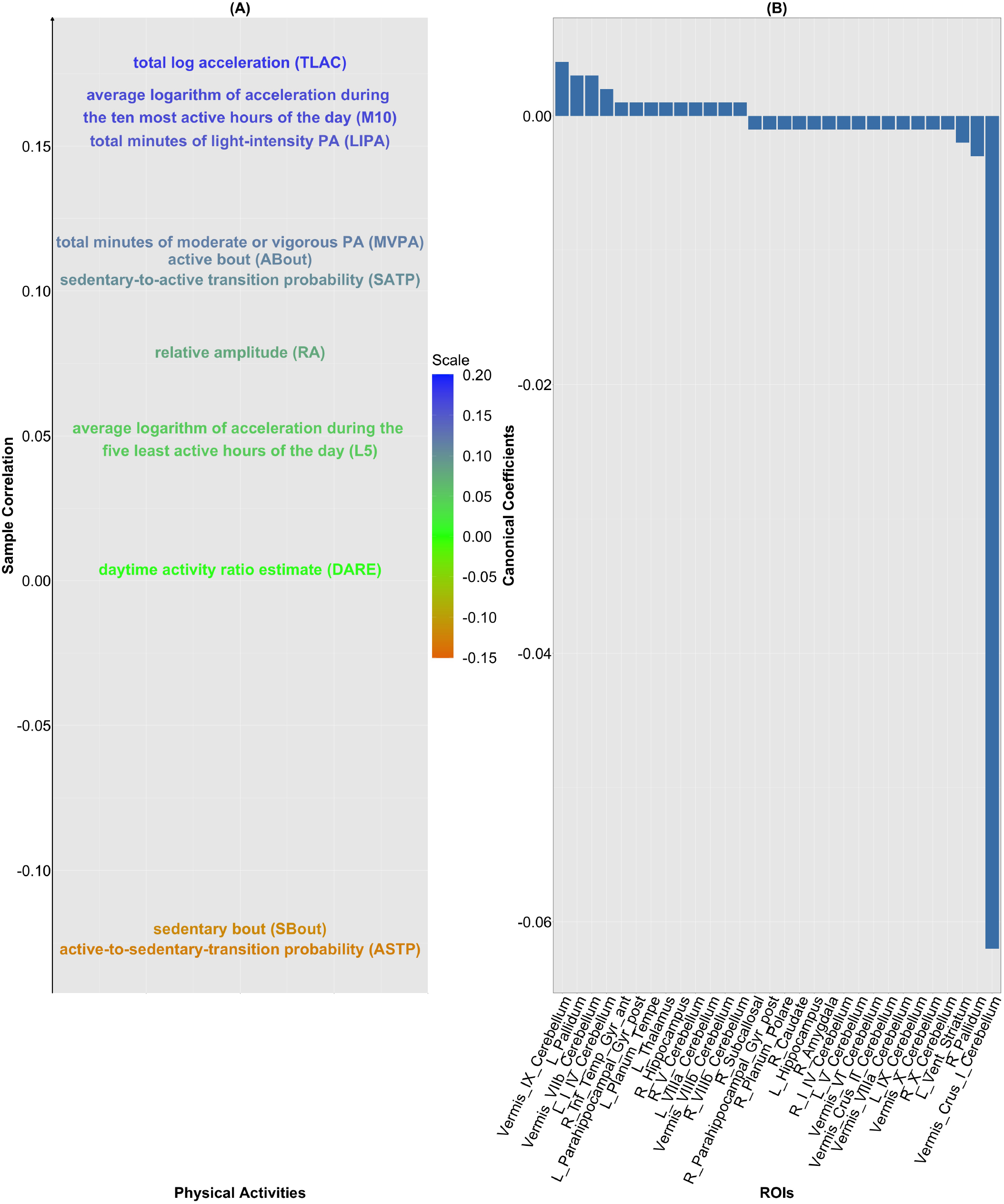
CCA relating GMV to PA variables. (A) The sample correlations between each residualized PA variable (each column of 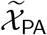) and the canonical variate 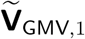; (B) the ROIs corresponding to the 30 largest elements in absolute scale of 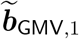.

#### Connection between PA variables and the Canonical Variate 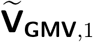

Sample correlations between each of the 11 residualized PA variables and the canonical variate 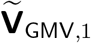 are shown in Figure 4(A). Four PA variables (TLAC, M10, LIPA and MVPA) that are deemed to reflect vigorous activity are positively correlated with 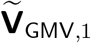, where the sample correlations are 0.18, 0.16, 0.15 and 0.12. In comparison, SBout, which measures inactivity is negatively correlated (sample correlation: -0.12) with 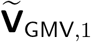. This leads to the conclusion that dynamic activity patterns positively correlate with GMV.

### 3.3 Variable Importance: Functional Connectome and Physical Activity

Let the mean-centered Fisher transformed FC matrix, confounders, and PA variables be **Y**_FC_, **G**_FC_ and **X**_PA_, respectively. Moreover, let **X**_(FC,PA)_ be the column-wise combination of **G**_FC_ and **X**_PA_, followed by demeaning and variance normalization. The statistical operations discussed in Section 2.4 are then applied to **Y**_FC_ as the matrix of response vectors and **X**_(FC,PA)_ as the design matrix.

The PA variables considered here are correlated and may reflect a small number of latent behavioral dimensions (e.g., overall activity intensity and circadian patterning). Consequently, variable importance results should be interpreted as identifying representative PA features rather than mechanistically distinct predictors.

#### Test of Significance of Linear Modeling and Prediction Assessment

Linear regression models are statistically significant in estimating 981 out of the 1485 columns (66.6%) of **Y**_FC_ by **X**_(FC,PA)_; the logarithm of the associated *p*-values after controlling for multiplicity are shown in Figure 5(A). Let the index set of these edges be *𝒥* _(FC,PA)_. To study the distribution of the significant connections, based on Figure 5(A), we find that 115 (11.7%), 41 (4.2%), 40 (4.08%) of the 981 edges are intra-region connections within the control and attention, cingulo-opercular and sensorimotor networks. Moreover, based on these connections in 𝒥_(FC,PA)_, predictive metrics in terms of the sample correlations (range: [-0.016, 0.25]) between observed and predicted functional connectomes are shown in Figure 5(B). It is clear that the intra-region edges in the sensorimotor network are better predicted by PA variables than the other networks due to the densely colored pattern. The average predictive metric for the edges in the sensorimotor network is 0.11, the highest among all five networks. Further, for 270 out of the 981 edges this metric exceeds 0.125. Among them, 77 (28.5%), 60 (22.2%) and 57 (21.1%) are from the control and attention, sensorimotor and cingulo-opercular networks, respectively. Hence, it appears that FC values facilitating the processing of the movement, especially those located in the sensorimotor and control and attention networks, are well predicted by the PA variables.

**Figure 5.**
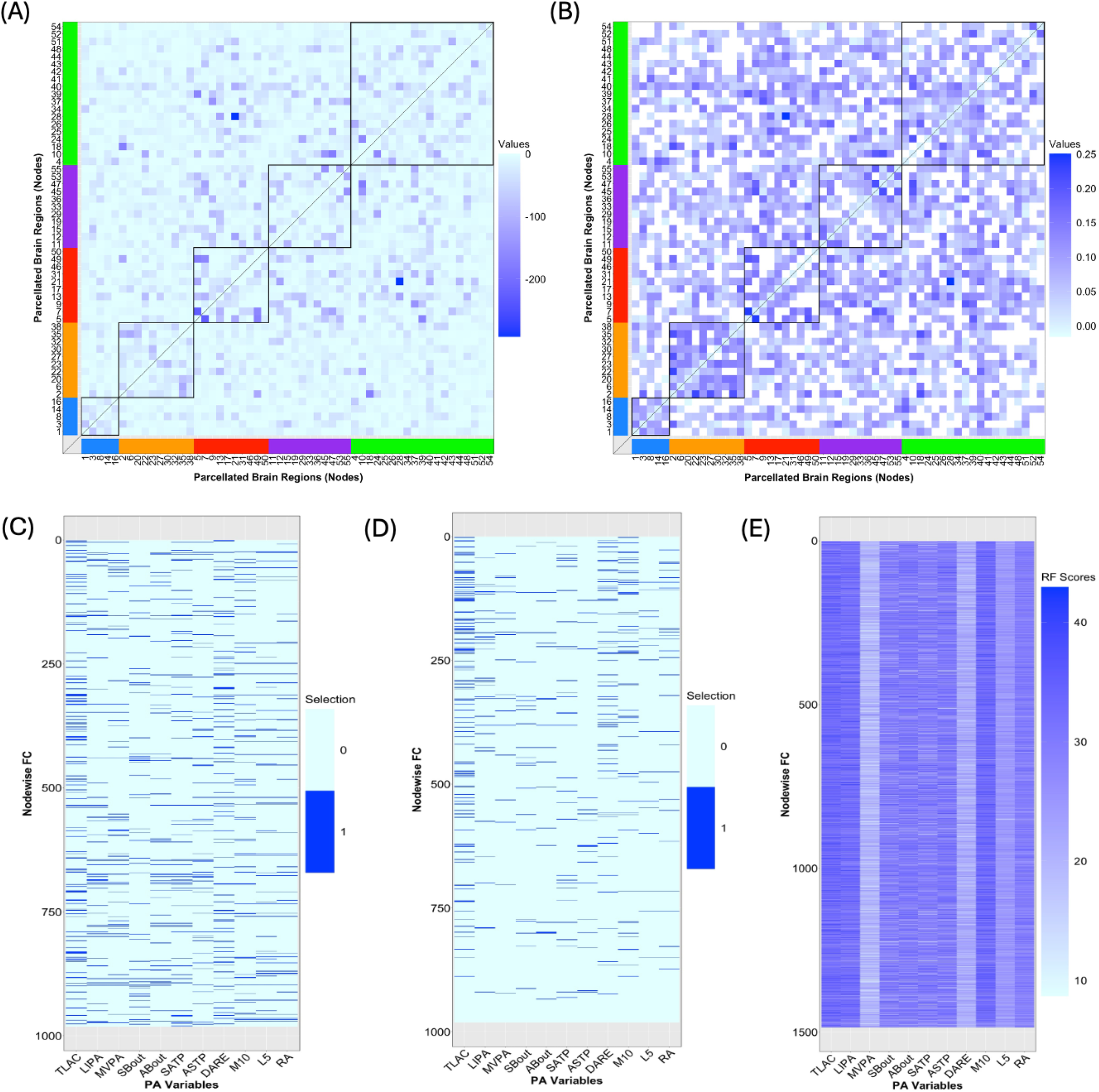
Prediction of FC by PA variables and assessment of variable importance of PA variables in modeling FC. (A) The logarithm of adjusted *p*-values associated with testing the significance of linear regression models when estimating FC by PA variables; the diagonal elements are omitted; (B) sample correlations between predicted and observed FCs corresponding to the locations identified in (A); (C) selection of PA variables determined by testing of individual regression coefficient by fitting of full linear regression model; (D) selection of PA variables via stepwise BIC; (E) significance of PA variables determined by random forest (RF).

#### Significance of PA Variables

Significance assessment of the PA variables from (*i*) fitting the full linear regression model, (*ii*) performing variable selection via stepwise BIC, and (*iii*) fitting random forest, is shown in Panels (C) to (E) in Figure 5. Clearly, variation exists in the results due to different modeling and selection mechanisms. However, a key feature shared across the three methods is that TLAC and M10 consistently rank among the most important PA variables. Moreover, the measure of importance of PA variables based on the scores obtained from the random forest (Panel (C)) closely aligns with the discovery from the sample correlations between each PA variable and the canonical variate for the functional connectomes 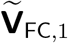 shown in Figure 2(C). This phenomenon highlights the crucial association of TLAC, M10 and LIPA, which quantify the volume, fragmentation and circadian rhythms aspects of physical activities, respectively, with functional connectivity.

### 3.4 Variable Importance: Gray Matter Volumes and Physical Activity

The inferential procedures in Section 2.4 are applied to **Y**_GMV_ as the matrix of response vectors and **X**_(GMV,PA)_ (consisting of confounders and PA variables) as the design matrix.

#### Test of Significance of Linear Modeling and Prediction Assessment

Linear regression models are statistically significant in modeling the GMVs associated with 136 out of the 139 ROIs (97.8%) by **X**_(GMV,PA)_; the logarithm of multiplicity-adjusted *p*-values is shown in Figures S12-S14 from Section 6 of the Supplementary Materials. Let the index set of these 136 ROIs be *𝒥* _(GMV,PA)_. Then, corresponding to ROIs belonging to *𝒥* _(GMV,PA)_, predictive metrics are obtained as the sample correlations between the observed and predicted GMVs and shown in Figure 6. The range of values is between 0.095 and 0.49, which is approximately twice as wide as that for the performance of the FC predicted by PA variables (Figure 5(B)). This indicates that the PA variables may be better predictors for GMV than FC. Moreover, the ROIs in *𝒥* _(GMV,PA)_ whose predictive metrics fall within the top 25^th^ percentile include prefrontal cortex, posterior cingulate gyrus, lingual gyrus, hippocampus and amygdala, which exhibit strong positive associations with physical activity.

**Figure 6.**
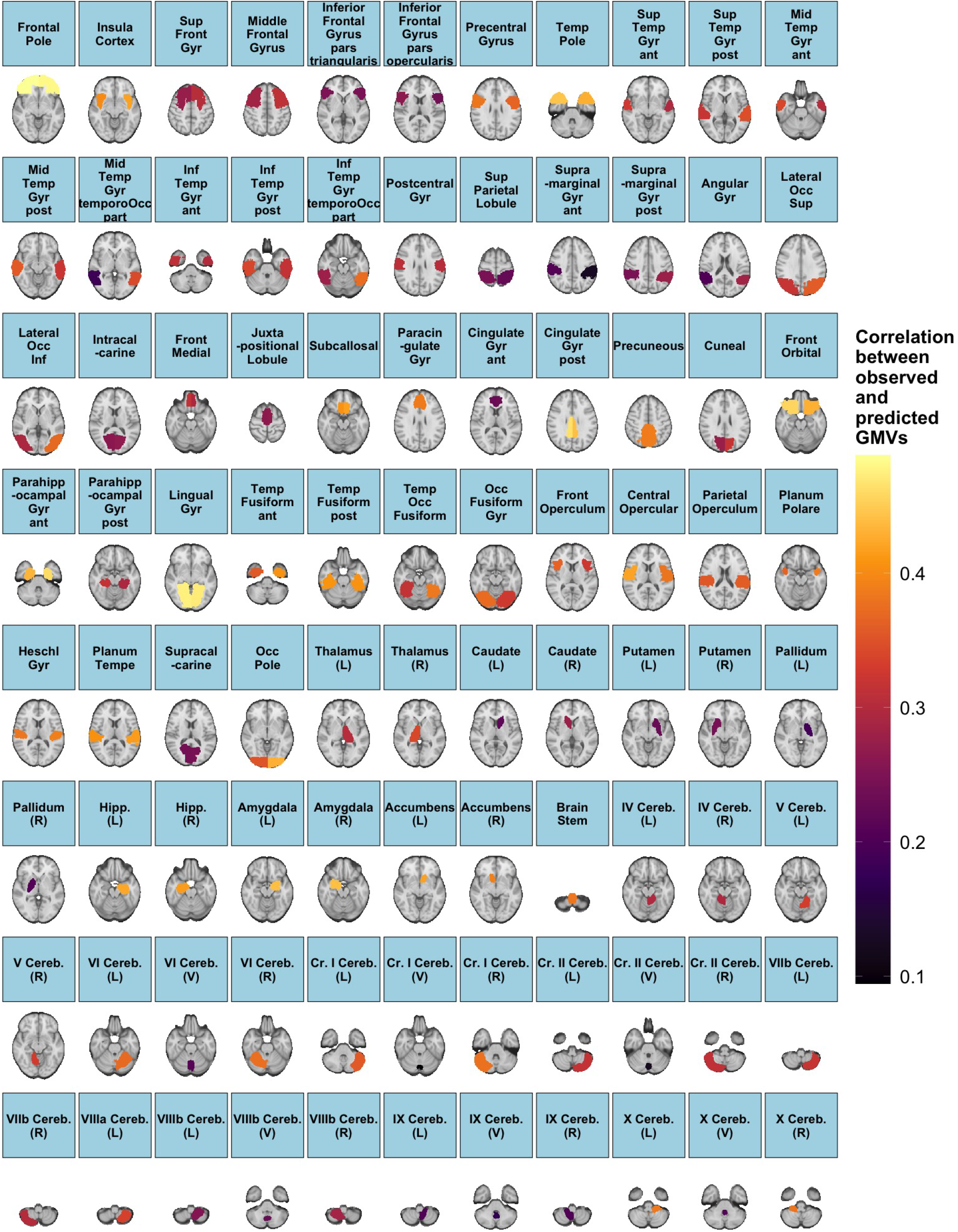
Prediction of GMVs by PA variables. The correlations between observed and predicted GMVs for 136 out of the 139 ROIs.

#### Significance of PA Variables

The variable importance of PA variables from (*i*) fitting the full linear regression model, (*ii*) performing variable selection via stepwise BIC, and (*iii*) fitting random forest is shown in Figure 7. Analogous to Figure 5(C)-(E), TLAC and M10 are identified among the key predictors based on the average selection score (column-wise) across the three approaches. In addition, from Figure 7(C), M10, TLAC, LIPA are the top three variables according to the scores assigned by the random forest. In fact, these three PA variables are most positively correlated with the canonical variate 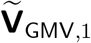 of the primary CCA mode (Figure 4(A)), further corroborating their importance in predicting GMV. On the other hand, as seen in Figure 7(C), variable importance of PA variables for GMVs corresponding to three groups of ROIs seem to deviate from the aforementioned pattern. These three clusters consist of 15 ROIs, which are the superior frontal gyrus, juxtapositional lobule cortex, paracingulate gyrus (left), paracingulate gyrus (right), anterior division (right) of cingulate gyrus, posterior division (right) of cingulate gyrus, caudate (left), caudate (right), putamen (left), putamen (right), pallidum (left), pallidum (right), brain-stem, crus I cerebellum (vermis) and VIIIb cerebellum (vermis). In fact, most of these ROIs have limited association with movement control. For example, superior frontal gyrus is involved in self-awareness, paracingulate gyrus is related to cognitive and affective regulation, cingulate gyrus facilitates the regulation of emotions and pain, caudate is important for learning, memory, reward, motivation, emotion, and romantic interaction, and pallidum helps process and execute motivated behaviors. As such, PA variables may not serve as proper predictors for GMV in these 15 ROIs.

**Figure 7.**
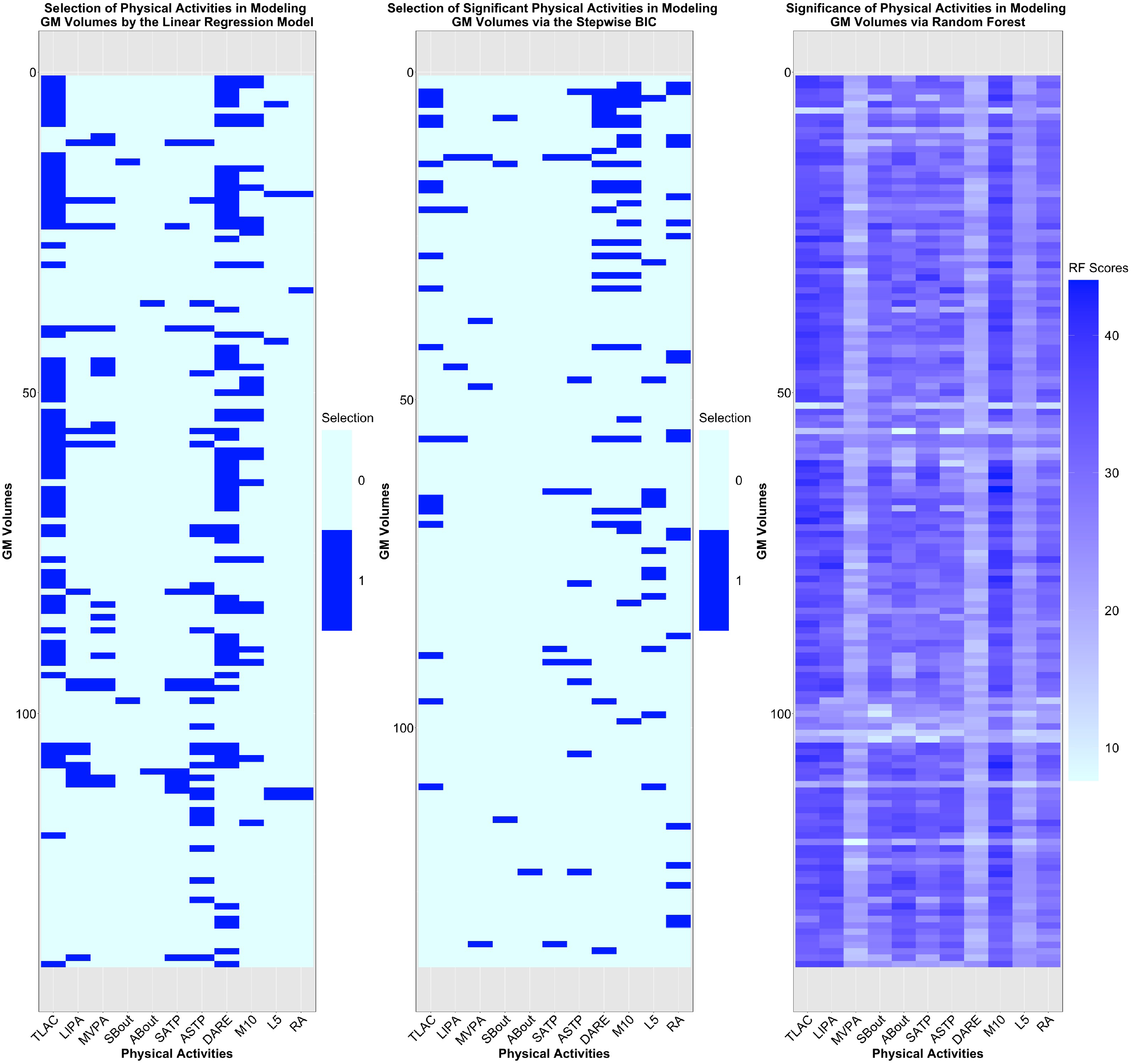
Assessment of importance of PA variables in modeling GMV. (A) Selection of PA variables determined by testing individual regression coefficient when fitting the full linear regression model; (B) selection of PA variables via stepwise BIC; (C) significance of PA variables determined by random forest (RF).

### 3.5 Risk Prediction: Functional Connectome and Physical Activity

Three logistic regression models (*M*_*i*_, *i* = 1, 2 and 3) were fit. Here **y**_(FC,PA,disease)_ denotes the 0-1 binary disease status, and **X**_(FC,PA)_(*M*_*i*_) the design matrix for the model *M*_*i*_, *i* = 1, 2 and 3. In particular, *M*_1_ denotes the full model containing PA variables and FC, *M*_2_ and *M*_3_ serve as submodels containing only PA variables and FC respectively.

To assess and compare the classification accuracy, the ROC curves and their associated AUCs corresponding to the nested logistic regression models are shown in Figure 8(A)-(D). In terms of AUC, we first observe that the accuracy for classifying the status of diabetes and CHD surpasses that for stroke and cancer across all the three nested models. The AUCs associated with these three models in the classification of CHD and diabetes status are approximately 0.8, while those for stroke and cancer are approximately 0.70 and 0.64. This underscores the stronger capability of the nested logistic regression models consisting of PA variables and FC in distinguishing the dichotomous disease status for CHD, diabetes, and stroke compared to cancer.

**Figure 8.**
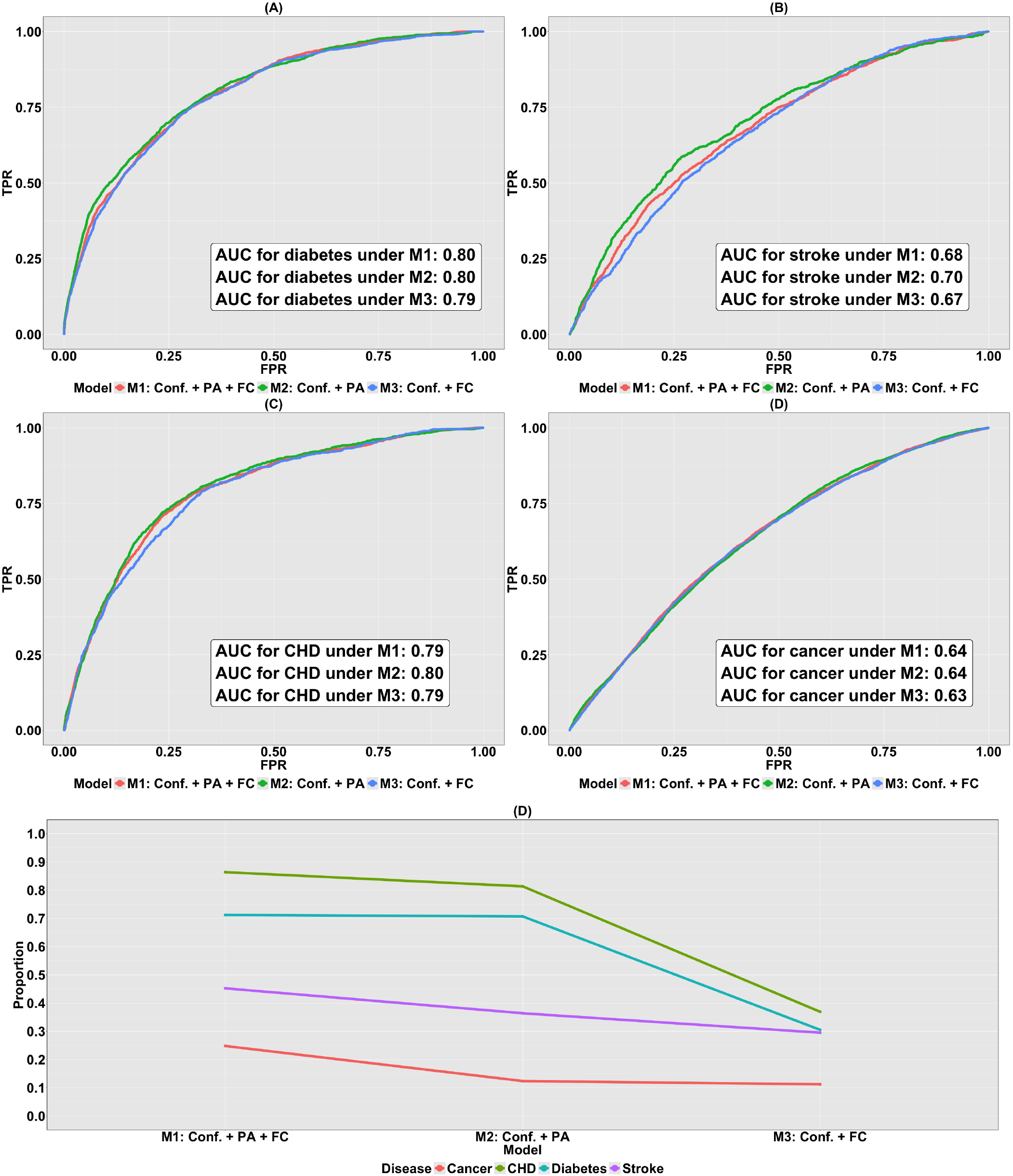
Disease risk prediction using FC and PA variables. (A)-(D) ROC curves for three nested logistic regression models (*M*_1_ to *M*_3_) fit to data consisting of FC and PA variables for classifying status of diabetes, stroke, CHD and cancer, respectively; (E) the proportion of variation in the binary disease status explained by *M*_1_, *M*_2_ and *M*_3_ fitted to data consisting of confounders, FC and PA variables. Here, *M*_1_ contains confounders, PA variables and FC. *M* 2 contains confounders and PA variables, and *M*_3_ contains confounders and FC.

Based on Figure 8(A)-(D), across all the four diseases, no notable discrepancies in the AUCs are observed among the three logistic models. In other words, *M*_1_, *M*_2_ and *M*_3_ demonstrate analogous predictive performances on classifying the status of each disease. Considering the limitations of the ROC-AUC method explained in Section 2.5, this particular assessment metric may not be informative and reliable due to the existence of highly imbalanced 0-1 binary disease status datasets, necessitating an alternative evaluation approach to ascertain the relative contribution of PA variables and FC to the variation in the data.

We calculate McFadden’s pseudo *R*^2^ (McFadden, 1974) for all the three logistic regression models illustrated in Figure 8(E). This metric shows the proportion of variation in the binary status of disease that can be explained by the model *M*. From Figure 8(E) two key observations are made. First, for logistic regression model *M*_*i*_, the benchmarks for the four diseases ranked in decreasing order are given by 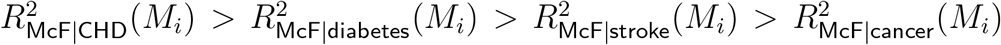. Hence the largest decline between two successive metrics exists between those computed for diabetes and stroke, and stroke and cancer. To be specific, 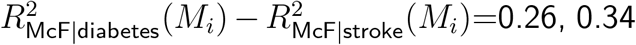 and 0.01, respectively, for *i* = 1, 2 and 3. In comparison, the counterparts for 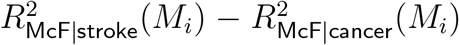 are 0.20, 0.24 and 0.18. This implies that CHD and diabetes can be uniformly better predicted by the logistic regression models consisting of PA variables and (or) FC, and the prediction considerably outperforms that for stroke and cancer. Second, for any disease in our study, 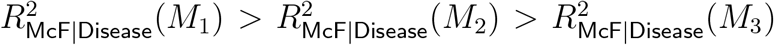.To be concrete, 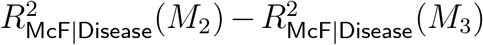 for CHD, diabetes, stroke and cancer are 0.44, 0.40, 0.069 and 0.012. These results highlight that PA variables consistently demonstrate greater predictive utility for cardiometabolic disease status than FC, suggesting limited incremental value of cross-sectional neuroimaging measures beyond behavioral indicators in this context.

### 3.6 Risk Prediction: Gray Matter Volumes and Physical Activity

The three logistic regression models (*M*_*i*_, *i* = 1, 2 and 3) were fit to the data consisting of the 0-1 binary response vector **y**_(GMV,PA,disease)_ and the design matrix **X**_(GMV,PA)_(*M*_*i*_). Here, *M*_1_ denotes the full model containing PA variables and GMVs, while *M*_2_ and *M*_3_ are the reduced models containing only PA variables and GMVs, respectively.

The ability of *M*_1_ to *M*_3_ to categorize the status of diabetes, stroke, CHD and cancer is shown by the ROC curves in Figure 9(A)-(D). Similar to when the brain IDP was FC, the performance of classifying disease status for CHD and diabetes exceeds that of stroke and cancer across all three models. Specifically, the associated AUCs for CHD, diabetes, stroke and cancer are in the close range, 0.81, 0.81, 0.70 and 0.63, respectively. Yet, it is ambiguous whether this pattern is attributable to PA variables or GMVs since for all diseases, *M*_1_ to *M*_3_ demonstrate almost the same classification accuracy. In addition, the presence of the imbalanced 0-1 binary disease status further raises concern on the validity of the ROC-AUC approach. To address the issue, we compute the McFadden’s pseudo *R*^2^ (McFadden, 1974).

**Figure 9.**
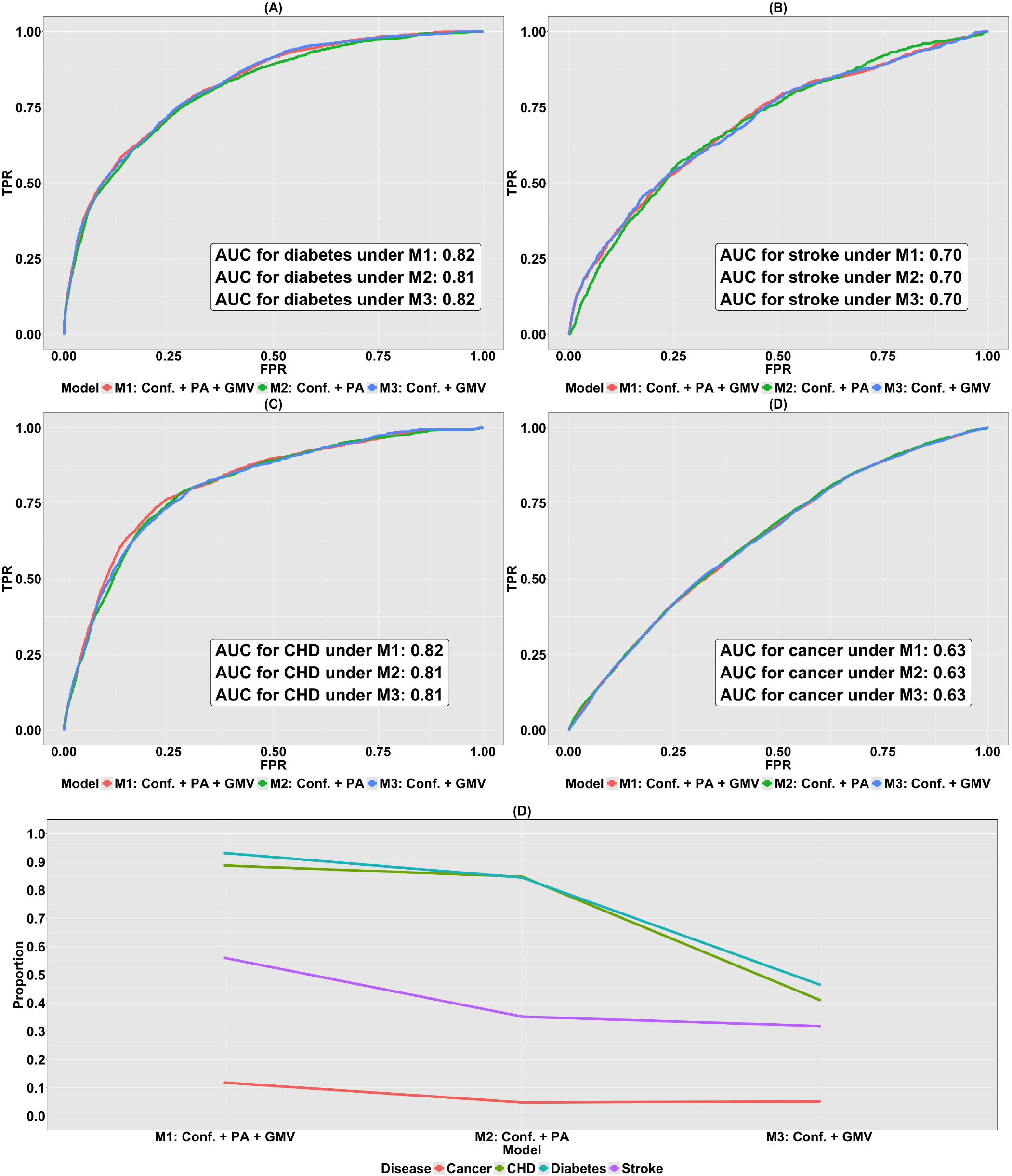
Disease risk prediction using GMV and PA variables. (A)-(D) ROC curves for three nested logistic regression models (*M*_1_ to *M*_3_) fit to data consisting of GMV and PA variables for classifying status of diabetes, stroke, CHD and cancer, respectively; (E) the proportion of variation in the binary disease status explained by *M*_1_, *M*_2_ and *M*_3_ fitted to data consisting of confounders, GMV and PA variables. Here, *M*_1_ consists of confounders, PA variables and GMV. *M* 2 contains confounders and PA variables, and *M*_3_ contains confounders and GMV.

Based on Figure 9(E), the values of this assessment metric are similar to when the brain IDP was FC: (*i*) for any model *M*_*i*_,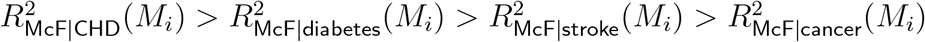, where considerable gaps have been observed to be between 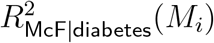 and 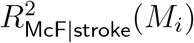, and 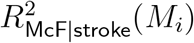 and 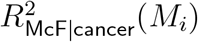, (*ii*) for any disease, 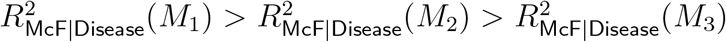. Both phenomena underscore the importance of PA variables in modeling and predicting disease status, especially for CHD and diabetes; the same assertion made in Section 3.5.

## 4 Discussion

This paper provides a systematic investigation of the links between brain imaging phenotypes and physical activity measurements, and their respective roles in disease modeling and prediction in the UK Biobank. Specifically, our work provides insights into six issues that, to date, have not been previously addressed in the literature, namely (*i*) the association between FC and PA variables, (*ii*) the association between GMV and PA variables, (*iii*) the significance of individual PA variables in estimating and predicting FC, (*iv*) the significance of individual PA variables in estimating and predicting GMV, (*v*) the relative contribution of PA and FC variables in predicting disease status, and (*vi*) the relative contribution of PA and GMV variables in predicting disease status. The first two issues are investigated using bipartial CCA, which allows one to evaluate the association between two multivariate datasets after removing the effects of confounder variables. The next two issues are addressed using a sequence of models, variable selection and machine learning procedures after accounting for the impacts of confounders. Finally, the last two issues are analyzed based on three nested logistic regression models, where the predictive and classification outcomes are measured by the ROC-AUC approach and the McFadden’s pseudo *R*^2^. Below follow some of the key conclusions of our analysis. Note that while the present analyses identify robust statistical associations between physical activity and brain imaging phenotypes, several limitations constrain interpretation. First, PA and imaging data were collected at different time points, limiting certain types of mechanistic inference. Second, CCA identifies shared low-dimensional structure but does not imply that the majority of brain variance is related to PA. These considerations underscore that the present findings are best interpreted as descriptive and predictive rather than mechanistic.

### Canonical Correlation Analysis of Brain IDPs and Physical Activity

We performed CCA to investigate the multivariate relationship between PA variables and two sets of IDPs, namely FC and GMV. For the CCA relating FC to PA variables, three CCA modes of population co-variation were found to be statistically significant based on both approximation and permutation tests. Focusing on the principal mode, the analysis revealed a high canonical correlation (0.5; see Figure 1(A)) and the strong linear association between the primary canonical variates (adjusted *R*^2^ : 0.25; see Figure 1(E)), indicating a statistically robust axis of shared variation between FC and PA.

Examination of the canonical coefficients for FC showed that the connections most strongly contributing to the primary canonical variate were primarily located within the control and attention networks, as well as the sensorimotor networks. Notably, four PA variables: total log acceleration (TLAC), light-intensity physical activity (LIPA), sedentary-to-active transition probability (SATP), and M10 (mean activity during the most active 10-hour window) were negatively associated with the primary FC canonical variate. In contrast, SBout, a measure of sedentary intensity, showed a positive association.

In the CCA between GMV and PA variables, four CCA modes were statistically significant based on the tests described above. However, concerns regarding the validity of the tests (see Section 2.3), the relatively low primary canonical correlation (0.19; see Figure 3(A)), and a weak linear association between the primary canonical variates (adjusted *R*^2^ : 0.04; see Figure 3(E)) suggest that caution should be taken in making interpretations, particularly beyond the primary mode. In contrast to the FC results, the GMV primary canonical variate was positively associated with four PA variables TLAC, M10, LIPA and moderate-to-vigorous physical activity (MVPA), and negatively associated with SBout.

### Predictive Modeling of Brain IDPs Using Physical Activity Variables

In modeling FC using PA variables, 66.6% of the 1485 nodewise functional connections were significantly predicted by linear regression models (Figure 5(A)). Moreover, higher predictive performance, measured by the correlation between observed and predicted values, was observed for connections within the control, attention, sensorimotor, and cingulo-opercular networks, with correlation values ranging from -0.016 to 0.25 (Figure 5(B)). In comparison, 97.8% of regional GMVs were significantly estimated by linear regression models incorporating PA variables (Supplementary Figures S12-S14), with predictive correlations ranging from 0.095 to 0.49 (Figure 6). Of particular note, ROIs such as the frontal pole (Erickson et al., 2014), posterior cingulate gyrus (Gogniat et al., 2022), lingual gyrus (Herting et al., 2016), hippocampus (Firth et al., 2018) and amygdala (Estévez-López et al., 2023) demonstrated strong positive association with physical activity, providing complementary quantitative evidence of the positive association between volumetric characteristics of isolated brain regions and physical activity.

### Variable Importance and Disease Risk Prediction

In quantifying the importance of individual PA variables in explaining variance in FC and GMV, three PA variables TLAC, M10 and LIPA were consistently identified as top contributors. These variables refer to total log acceleration, average log acceleration during the most active 10 hours, and light intensity physical activity, respectively. These variables are correlated and may reflect a small number of latent behavioral dimensions, including overall activity intensity and circadian rhythms (Leroux et al., 2020).

Finally, we assessed the predictive utility of brain IDPs and PA variables for four health conditions: CHD, diabetes, stroke, and cancer. Using nested logistic regression models, we found classification performance, measured using the area under the ROC curve, was highest for CHD and diabetes for both FC (Figure 8(A)-(D)) and GMV (Figure 9(A)-(D)). Computation of McFadden’s pseudo *R*^2^ for the three models showed that PA variables alone contributed the most predictive value across all four disease types (Figure 8(E) and Figure 9(E)), suggesting that cross-sectional neuroimaging measures provide limited incremental explanatory value beyond behavioral indicators in this context. In contrast, the relatively weak predictive performance of FC or GMV alone may reflect their indirect relationship with these disease outcomes. Previous studies have shown that these diseases are associated with disruptions in FC and alterations in GMV (Dai et al., 2024; Hall et al., 2021; Vuorinen et al., 2014; Falcó-Roget et al., 2024). However, reverse modeling and prediction of the diseases by brain IDPs does not establish a reliable, strong predictive performance, leading to the low 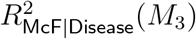 (*M*_3_) observed in Panel (E) of both Figures 8 and 9. Focusing on the performance of PA variables, our findings corroborate existing scientific findings, indicating that physical activity is foundational in improving blood sugar control and increasing insulin sensitivity, thereby mitigating the risks of CHD, diabetes and stroke. On the other hand, with regard to the prediction of cancer using PA variables, factors contributing to cancer risks are multifaceted, including but not limited to genetic mutations, lifestyle choices and environmental influences. Moreover, in our study, cancer type was not specified. For these reasons, while physical activity is shown to be associated with certain types of cancer including bladder cancer (Moore et al., 2016), breast cancer (Pizot et al., 2016), colon cancer (Liu et al., 2016) and kidney (renal cell) cancer (Behrens & Leitzmann, 2013), studies have also shown that physical activities is not consistently associated with risks of other cancer types, for example the subtype-specific hematologic cancers (Jochem et al., 2014). Therefore,the 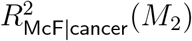 is low in this case.

### Conclusions

This study used CCA to examine associations between PA and two sets of brain IDPs (FC and GMV). The primary CCA mode showed a stronger association between FC and PA (canonical correlation = 0.50) than between GMV and PA (0.19), with key contributions from motor and attention networks. Notably, these associations were low-dimensional with respect to the brain IDPs, indicating that PA covaries with a relatively narrow subspace of FC and GMV variation. Vigorous PA measures (e.g., TLAC, LIPA, M10) were negatively associated with FC but positively associated with GMV, while sedentary behavior showed opposite patterns. Predictive modeling demonstrated that PA variables significantly predicted both FC and GMV, and generally outperformed brain IDPs in predicting disease risk, especially for diabetes and CHD. Three PA variables (TLAC, M10, and LIPA) consistently emerged as key contributors, although these predictors are correlated and may reflect a small number of latent behavioral dimensions (e.g., activity intensity and circadian rhythm). Overall, the findings support robust descriptive and predictive relationships between physical activity, brain structure, brain function, and disease status.

## Data and Code Availability

Code to reproduce analyses is available on GitHub: https://github.com/Dongliang-JHU/PA-Brain-IDPs-UKB. Analyses relied on the UK Biobank.

## Author Contributions

Dongliang Zhang: Conceptualization, Methodology, Software, Validation, Formal Analysis, Investigation, Resources, Data Curation, Writing—Original Draft, Writing—Review & Editing, Visualization. Andrew Leroux: Writing-Review & Editing, Supervision. Ciprian M. Crainiceanu: Writing-Review & Editing, Supervision. Martin A. Lindquist: Conceptualization, Methodology, Validation, Formal Analysis, Resources, Writing—Original Draft, Writing—Review & Editing, Supervision, Project Administration, Funding Acquisition.

## Funding

This work was supported in part by NIH grant R01 EB026549 from the National Institute of Biomedical Imaging and Bioengineering (NIBIB), and R01MH129397 from NIMH.

## Declaration of Competing Interests

The authors declare that they have no known competing financial interests or personal relationships that could have appeared to influence the work reported in this paper.

## Acknowledgements

We are grateful to UK Biobank and the UK Biobank participants for making the resource data possible, and to the data processing team at Oxford University for sharing the processed data. The UK Biobank imaging project is funded by the Medical Research Council and the Wellcome Trust.

## Supplementary Material

Supplementary Material is available on the Github: https://github.com/Dongliang-JHU/PA-Brain-IDPs-UKB.

## Notes

### Competing Interest Statement

The authors have declared no competing interest.

