## Supplementary Materials for "Exploring Links between Brain Image-Derived Phenotypes and Accelerometer-Measured Physical Activity in the UK Biobank"

**Contents**

1 **Population Characteristics for Some Quantities in CCA** 1

2 **Population Characteristics for Some Quantities in Variable Importance Assessment** 4

3 **Graphical Representation of Brain Regions** 6

4 **Flow Charts for Data Processing** 10

5 **Population-level Pattern of the Functional Connectivity for CCA and Variable Importance Assessment** 17

6 **Testing of Significance of Linear Regression Models in Estimating the GMVs of the 139 ROIs** 18

**7 Method for Canonical Correlation Analysis (CCA) 21**

### List of Tables

S1 Population characteristics of the confounder variables for FC and GMV in CCA. For categorical variables, size and percentage of each category is shown. For continuous variables, mean (standard deviation (SD)) followed by (minimum, median, maximum and inter-quantile range (IQR)) is shown. 1

S2 Population characteristics of the confounder variables for PA in CCA. For categorical variables, size and percentage of each category is shown. For continuous variables, mean (standard deviation (SD)) followed by (minimum, median, maximum and inter-quantile range (IQR)) is shown. . . . . 2

S3 Population characteristics for PA variables in CCA when respectively relating FC and GMV to PA variables. Mean (standard deviation (SD)) followed by (minimum, median, maximum and inter-quantile range (IQR)) is shown. . . . . . . . . . . . . . . . . . . . . . . . . . . . . . . . . . . 3

S4 Population characteristics of confounders for assessing variable importance (VI) of PA variables in modeling FC and GMV. For categorical variables, size and percentage of each category is shown. For continuous variables, mean (standard deviation (SD)) followed by (minimum, median, maximum and inter-quantile range (IQR)) is shown. . . . . . . . . . . . . . . . . . . . . 4

S5 Population characteristics of PA variables for accessing their variable importance in modeling FC and GMV. Mean (standard deviation (SD)) followed by (minimum, median, maximum and inter-quantile range (IQR)) is shown. . . . . . . . . . . . . . . . . . . . . . . . . . . . . . . . . . . . 5

### List of Figures

S1 The 96 ROIs from the Harvard-Oxford cortical structural atlas. . . . . . . 6
S2 The 15 ROIs from the Harvard-Oxford subcortical structural atlas. . . . . 7
S3 The 28 ROIs from the Diedrichsen structural atlas. . . . . . . . . . . . . . 8
S4 The 55 non-artefact components from parcellations of the brain obtained from the rs-fMRI pipeline in UKB. . . . . . . . . . . . . . . . . . . . . . . . 9
S5 Data processing flow chart of CCA relating FC to PA variables. . . . . . . 11
S6 Data processing flow chart of CCA relating GMV to PA variables. . . . . . 12
S7 Data processing flow chart for assessing variable importance (VI) of individual PA variable in modeling FC. . . . . . . . . . . . . . . . . . . . . . 13
S8 Data processing flow chart for assessing variable importance (VI) of individual PA variable in modeling GMV. . . . . . . . . . . . . . . . . . . . . . . 14
S9 Data processing flow chart for predicting the risks of diabetes, stroke, CHD and cancer via nested logistic models containing FC and PA. . . . . . . . . 15
S10 Data processing flow chart for predicting the risks of diabetes, stroke, CHD and cancer by nested logistic models containing GMV and PA. . . . . . . . 16
S11 (A) mean functional connectivity matrix used in CCA relating FC to PA variables; (B) mean functional connectivity matrix used in assessing the importance of individual PA variable when modeling FC. The parcellated brain regions (nodes) are grouped by the five resting-state networks (RSNs). Specifically, the blue, orange, red, purple and green networks respectively controls visual, sensorimotor, default, cingulo-opercular, and control and attention functions. The diagonal elements are omitted. . . . . . . . . . . . 17
S12 The logarithm of adjusted $p$-values associated with testing the significance of linear regression models in estimating the GMVs of the 96 ROIs from the Harvard-Oxford cortical structural atlas. . . . . . . . . . . . . . . . . . . 18
S13 The logarithm of adjusted $p$-values associated with testing the significance of linear regression models in estimating the GMVs of the 15 ROIs from the Harvard-Oxford subcortical structural atlas. . . . . . . . . . . . . . . . . 19
S14 The logarithm of adjusted $p$-values associated with testing the significance of linear regression models in estimating the GMVs of the 28 ROIs from the Diedrichsen structural atlas. . . . . . . . . . . . . . . . . . . . . . . . . . 20

### 1. Population Characteristics for Some Quantities in CCA

*Table S1*

*Population characteristics of the confounder variables for FC and GMV in CCA. For categorical variables, size and percentage of each category is shown. For continuous variables, mean (standard deviation (SD)) followed by (minimum, median, maximum and inter-quantile range (IQR)) is shown.*

| Variables | UKB ID | Measurement of Confounders for FC (N=8580) (CCA relating FC to PA) | Measurement of Confounders for GMV (N=9208) (CCA relating GMV to PA) |
|---|---|---|---|
| Sex | 31 | Female: 4743 (55.4%)<br>Male: 3837 (44.6%) | Female: 5066 (55.0%)<br>Male: 4142 (45.0%) |
| Age | 34,52,53 | 55.68 (7.47)<br>(40, 56, 70, 12) | 55.72 (7.50)<br>(40, 56, 70, 12) |
| Head size | 25000 | 1.30 (0.12)<br>(0.92, 1.30, 1.74, 0.17) | 1.30 (0.12)<br>(0.92, 1.30, 1.76, 0.17) |
| Brain position (lateral) | 25756 | 0.41 (2.67)<br>(-35.01, 0.28, 17.28, 3.25) | 0.41 (2.66)<br>(-35.01, 0.29, 17.28, 3.24) |
| Brain position (transverse) | 25757 | 63.69 (5.37)<br>(53, 62, 95, 7) | 63.60 (5.34)<br>(53, 62, 95, 7) |
| Brain position (longitudinal) | 25758 | -16.64 (32.04)<br>(-92.06, -27.65, 98.98, 51.84) | -16.69 (32.09)<br>(-92.06, -27.65, 98.98, 51.69) |
| Scanner table position | 25759 | -1066.79 (31.24)<br>(-1187, -1042, -1042, 52) | -1066.91 (31.24)<br>(-1187, -1042, -1036, 53) |
| Mean rfMRI head motion | 25741 | 0.12 (0.06)<br>(0.03, 0.11, 1.39, 0.06) | -<br>- |
| median absolute rfMRI head motion | 24439 | 0.65 (0.47)<br>(0.09, 0.52, 10.11, 0.38) | -<br>- |
| Head motion in T1 | 24419 | -<br>- | 0.31 (0.17)<br>(-0.15, 0.29, 6.96, 0.18) |
| Intensity scaling for rfMRI | 25929 | 1.96 (0.19)<br>(1, 2, 2, 0) | -<br>- |
| Intensity scaling for T1 | 25925 | -<br>- | 5.82 (0.94)<br>(1, 6, 6, 0) |

*Table S2*

*Population characteristics of the confounder variables for PA in CCA. For categorical variables, size and percentage of each category is shown. For continuous variables, mean (standard deviation (SD)) followed by (minimum, median, maximum and inter-quantile range (IQR)) is shown.*

| Variables | UKB ID | Measurement of Confounders for PA (N=8580) (CCA relating FC to PA) | Measurement of Confounders for PA (N=9208) (CCA relating GMV to PA) |
|---|---|---|---|
| Sex | 31 | Female: 4743 (55.4%)<br>Male: 3837 (44.6%) | Female: 5066 (55.0%)<br>Male: 4142 (45.0%) |
| Age | 34,52,53 | 55.68 (7.47)<br>(40, 56, 70, 12) | 55.72 (7.50)<br>(40, 56, 70, 12) |
| BMI | 21001 | 26.50 (4.26)<br>(15.94, 25.9, 56.6, 5.19) | 26.53 (4.31)<br>(15.94, 25.92, 63.58, 5.19) |
| Smoking | 20116 | Never: 5229 (60.9%)<br>Previous: 2845 (33.3%)<br>Current: 489 (5.7%)<br>Prefer not to answer: 17 (0.2%) | Never: 5597 (60.8%)<br>Previous: 3067 (33.3%)<br>Current: 525 (5.7%)<br>Prefer not to answer: 19 (0.2%) |
| Alcohol drinking | 1558 | Daily or almost daily: 1843 (21.5%)<br>Three or four times a week: 2421 (28.2%)<br>Once or twice a week: 2214 (25.8%)<br>One to three times a month: 946 (11.0%)<br>Special occasions only: 754 (8.8%)<br>Never: 401 (4.7%)<br>Prefer not to answer: 1 (0%) | Daily or almost daily: 1965 (21.3%))<br>Three or four times a week: 2604 (28.3%)<br>Once or twice a week: 2369 (25.7%)<br>One to three times a month: 1011 (11.0%)<br>Special occasions only: 818 (8.9%)<br>Never: 440 (4.8%)<br>Prefer not to answer: 1 (0%) |
| Long-standing illness | 2188 | Yes: 1951 (22.7%)<br>No: 6487 (75.6%)<br>Do not know: 138 (1.7%)<br>Prefer not to answer: 4 (0%) | Yes: 2120 (23.0%)<br>No: 6938 (75.3%)<br>Do not know: 145 (1.6%)<br>Prefer not to answer: 5 (0%) |
| Diabetes | 90010,130706, 130708,130710,130712 | Yes: 367 (4.3%)<br>No: 8213 (95.7%) | Yes: 414 (4.5%)<br>No: 8794 (95.5%) |
| Stroke | 90010,131180,131366,131368, 131362,131360,131056 | Yes: 194 (2.3%)<br>No: 8386 (97.7%) | Yes: 201 (2.2%)<br>No: 9007 (97.8%) |
| CHD | 90010,131306 | Yes: 269 (3.1%)<br>No: 8311 (96.9%) | Yes: 302 (3.3%)<br>No: 8609 (96.7%) |
| Cancer | 90010, 40005 | Yes: 1063 (12.4%)<br>No: 7517 (87.6%) | Yes: 1149 (12.5%)<br>No: 8059 (87.5%) |

*Table S3*

*Population characteristics for PA variables in CCA when respectively relating FC and GMV to PA variables. Mean (standard deviation (SD)) followed by (minimum, median, maximum and inter-quantile range (IQR)) is shown.*

| Variables | PA Measurement $\mathbf{X}_{\text{PA}}$ (N=8580) (CCA relating FC to PA) | PA Measurement $\mathcal{X}_{\text{PA}}$ (N=9208) (CCA relating GMV to PA) |
|---|---|---|
| TLAC | 3440.84 (310.95) (2040.56, 3440.77, 4960.09, 409.08) | 3439.31 (311.25) (2040.56, 3438.75, 4960.09, 409.41) |
| LIPA | 386.15 (91.21) (79.75, 382.20, 799.33, 121.72) | 385.66 (91.31) (79.75, 381.63, 799.33, 122.00) |
| MVPA | 19.23 (17.81) (0.00, 14.08, 154.83, 19.70) | 19.14(17.88) (0.00, 14.00, 154.83, 19.67) |
| SBout | 14.81 (3.97) (4.64, 14.15, 57.40, 4.81) | 14.81 (3.97) (4.64, 14.15, 57.40, 4.83) |
| ABout | 5.64 (1.50) (2.04, 5.41, 16.25, 1.85) | 5.63 (1.50) (2.04, 5.39, 16.25, 1.85) |
| SATP | 0.08 (0.02) (0.03, 0.07, 0.23, 0.02) | 0.08 (0.02) (0.03, 0.07, 0.23, 0.02) |
| ASTP | 0.20 (0.05) (0.07, 0.20, 0.54, 0.06) | 0.20 (0.05) (0.07, 0.20, 0.54, 0.06) |
| DARE | 0.66 (0.03) (0.32, 0.66, 0.78, 0.04) | 0.66 (0.03) (0.32, 0.66, 0.78, 0.04) |
| M10 | 3.34 (0.32) (1.70, 3.35, 4.48, 0.43) | 3.34 (0.32) (1.70, 3.35, 4.48, 0.42) |
| L5 | 0.99 (0.11) (0.37, 0.98, 2.67, 0.12) | 0.99 (0.11) (0.37, 0.98, 2.67, 0.12) |
| RA | 0.54(0.05) (0.18, 0.55, 0.77, 0.06) | 0.54 (0.05) (0.18, 0.55, 0.77, 0.06) |

### 2. Population Characteristics for Some Quantities in Variable Importance Assessment

*Table S4*

*Population characteristics of confounders for assessing variable importance (VI) of PA variables in modeling FC and GMV. For categorical variables, size and percentage of each category is shown. For continuous variables, mean (standard deviation (SD)) followed by (minimum, median, maximum and inter-quantile range (IQR)) is shown.*

| Variables | UKB ID | Measurement of Confounders (N=8582) (Modeling FC by PA) | Measurement of Confounders (N=9217) (Modeling GMV by PA) |
|---|---|---|---|
| Sex | 31 | Female: 4744 (55.3%)<br>Male: 3838 (44.7 %) | Female: 5070 (55.0%)<br>Male: 4147 (45.0%) |
| Age | 34,52,53 | 55.68 (7.47)<br>(40, 56, 70, 12) | 55.72 (7.50)<br>(40, 56, 70, 12) |
| BMI | 21001 | 26.50 (4.25)<br>(15.94, 25.91, 56.6, 5.19) | 26.53 (4.31)<br>(15.94, 25.92, 63.58, 5.19) |
| Smoking | 20116 | Never: 5229 (60.9%)<br>Previous: 2847 (33.2%)<br>Current: 489 (5.7%)<br>Prefer not to answer: 17 (0.2%) | Never: 5604 (60.8%)<br>Previous: 3069 (33.3%)<br>Current: 525 (5.70%)<br>Prefer not to answer: 19 (0.2%) |
| Alcohol drinking | 1558 | Daily or almost daily: 1843 (21.5%)<br>Three or four times a week: 2423 (28.2%)<br>Once or twice a week: 2214 (25.8%)<br>One to three times a month: 946 (11.0%)<br>Special occasions only: 754 (8.8%)<br>Never: 401 (4.7%)<br>Prefer not to answer: 1 (0%) | Daily or almost daily: 1966 (21.3%))<br>Three or four times a week: 2608 (28.3%)<br>Once or twice a week: 2371 (25.7%)<br>One to three times a month: 1012 (11.0%)<br>Special occasions only: 819 (8.9%)<br>Never: 440 (4.8%)<br>Prefer not to answer: 1 (0%) |
| Long-standing illness | 2188 | Yes: 1952 (22.7%)<br>No: 6488 (75.6%)<br>Do not know: 138 (1.7%)<br>Prefer not to answer: 4 (0%) | Yes: 2123 (23.0%)<br>No: 6944 (75.3%)<br>Do not know: 145 (1.6%)<br>Prefer not to answer: 5 (0%) |
| Diabetes | 90010,130706, 130708,130710,130712 | Yes: 367 (4.3%)<br>No: 8215 (95.7%) | Yes: 415 (4.5%)<br>No: 8802 (95.5%) |
| Stroke | 90010,131180,131366,131368, 131362,131360,131056 | Yes: 194 (2.3%)<br>No: 8388 (97.7%) | Yes: 201 (2.2%)<br>No: 9016 (97.8%) |
| CHD | 90010,131306 | Yes: 269 (3.1%)<br>No: 8313 (96.9%) | Yes: 302 (3.3%)<br>No: 8915 (96.7%) |
| Cancer | 90010, 40005 | Yes: 1064 (12.4 %)<br>No: 7518 (87.6 %) | Yes: 1149 (12.5%)<br>No: 8068 (87.5%) |

*Table S5*

*Population characteristics of PA variables for accessing their variable importance in modeling FC and GMV. Mean (standard deviation (SD)) followed by (minimum, median, maximum and inter-quantile range (IQR)) is shown.*

| Variables | Measurement of PA in modeling FC (N=8582) | Measurement of PA in modeling GMV (N=9217) |
|---|---|---|
| TLAC | 3440.81 (310.92)<br>(2040.56, 3440.66, 4960.09, 409.02) | 3439.32 (311.22)<br>(2040.56, 3438.71, 4960.09, 409.37) |
| LIPA | 386.13 (91.22)<br>(79.75, 382.18, 799.33, 121.85) | 385.67 (91.31)<br>(79.75, 381.67, 799.33, 122.00) |
| MVPA | 19.23 (17.81)<br>(0.00, 14.00, 154.83, 19.70) | 19.13 (17.87)<br>(0.00, 14.00, 154.83, 19.67) |
| SBout | 14.81 (3.97)<br>(4.64, 14.15, 57.40, 4.81) | 14.81 (3.97)<br>(4.64, 14.15, 57.40, 4.83) |
| ABout | 5.64 (1.51)<br>(2.04, 5.41, 16.25, 1.85) | 5.63 (1.50)<br>(2.04, 5.39, 16.25, 1.85) |
| SATP | 0.08 (0.02)<br>(0.03, 0.07, 0.23, 0.02) | 0.08 (0.02)<br>(0.03, 0.07, 0.23, 0.02) |
| ASTP | 0.20 (0.05)<br>(0.07, 0.20, 0.54, 0.06) | 0.20 (0.05)<br>(0.07, 0.20, 0.54, 0.06) |
| DARE | 0.66 (0.03)<br>(0.32, 0.66, 0.78, 0.04) | 0.66 (0.03)<br>(0.32, 0.66, 0.78, 0.04) |
| M10 | 3.34 (0.32)<br>(1.70, 3.35, 4.48, 0.43) | 3.34 (0.32)<br>(1.70, 3.35, 4.48, 0.42) |
| L5 | 0.99 (0.11)<br>(0.37, 0.98, 2.67, 0.12) | 0.99 (0.11)<br>(0.37, 0.98, 2.67, 0.12) |
| RA | 0.54 (0.05)<br>(0.18, 0.55, 0.77, 0.06) | 0.54 (0.05)<br>(0.18, 0.55, 0.77, 0.06) |

### 3. Graphical Representation of Brain Regions

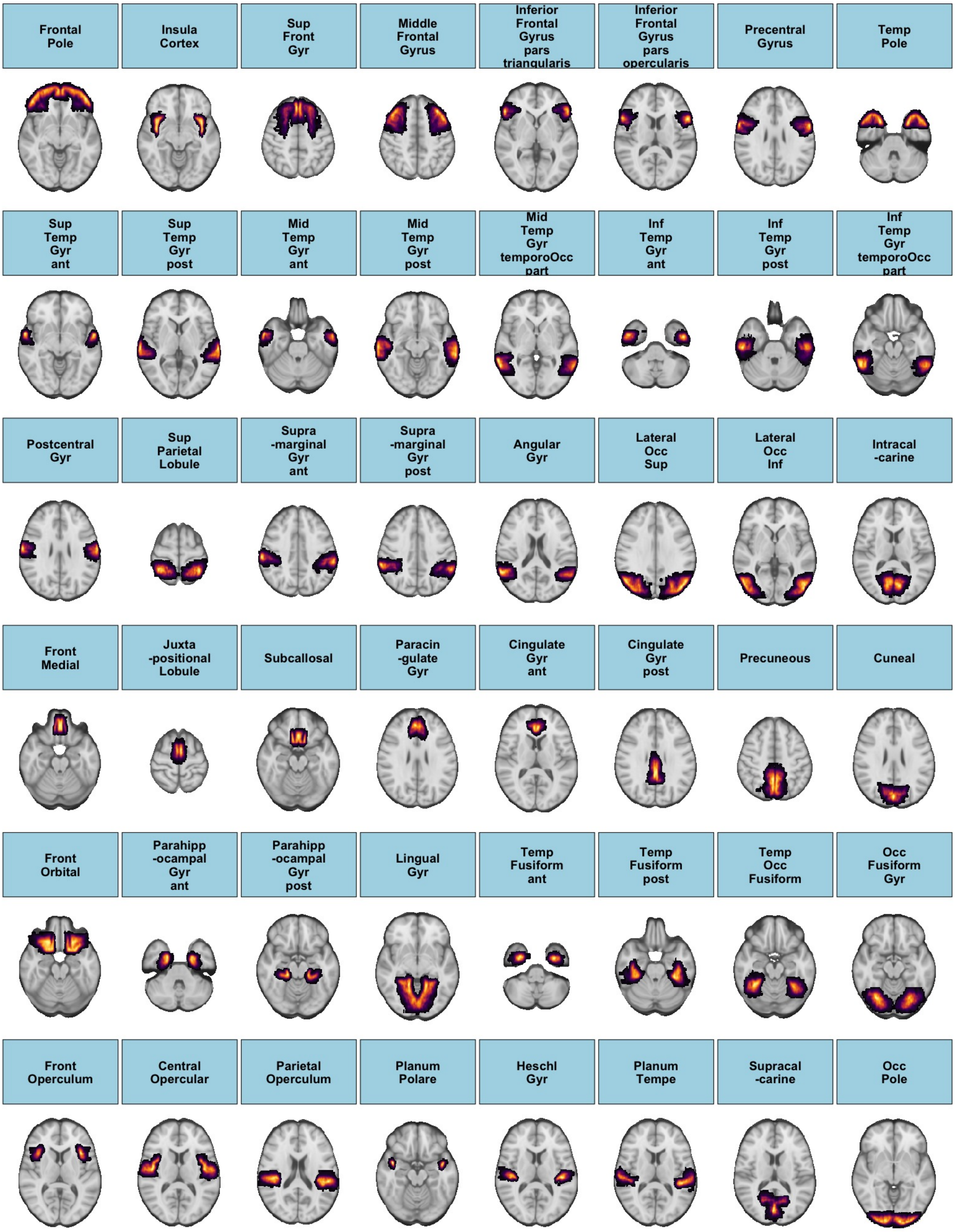


*Figure S1. The 96 ROIs from the Harvard-Oxford cortical structural atlas.*

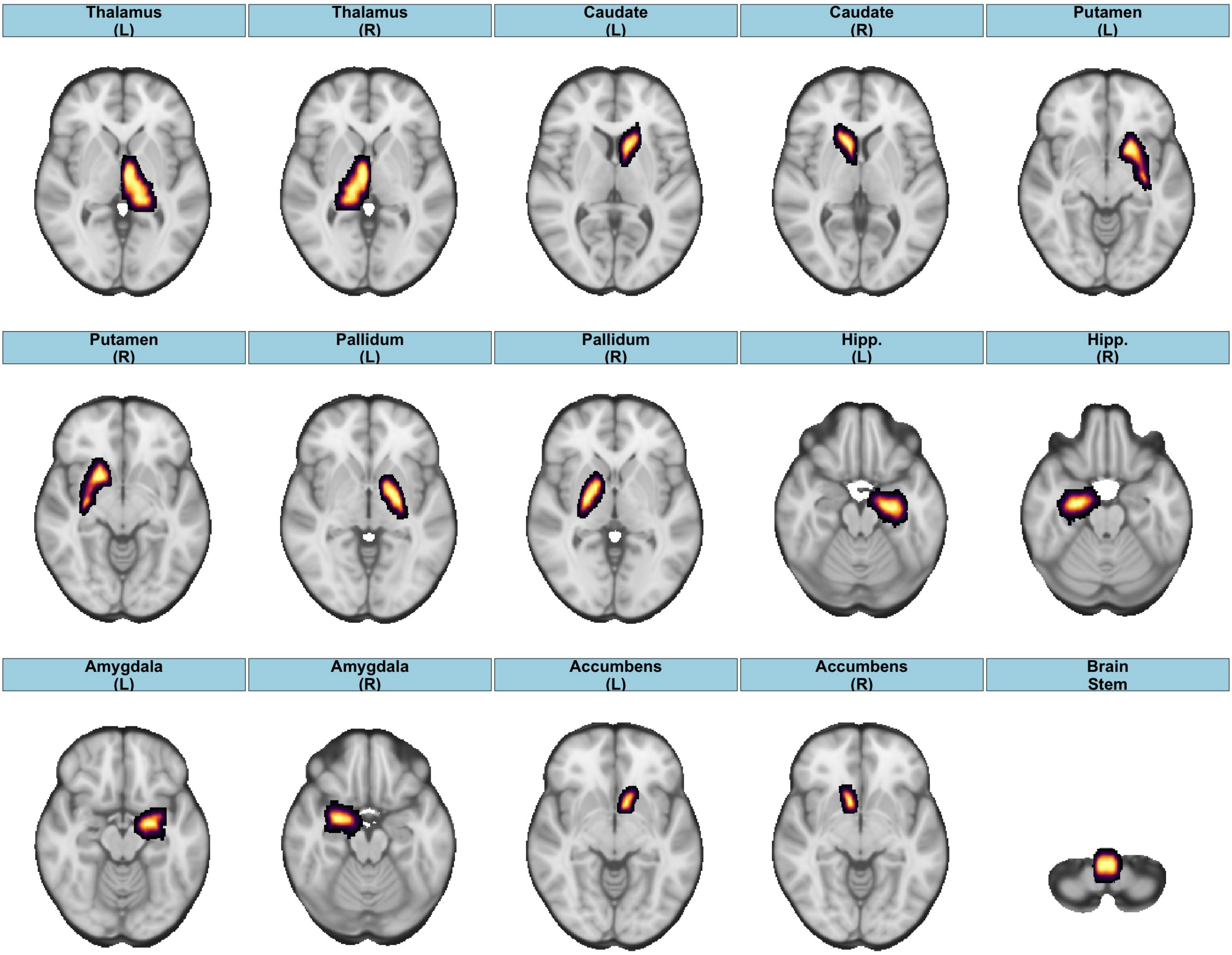


*Figure S2. The 15 ROIs from the Harvard-Oxford subcortical structural atlas.*

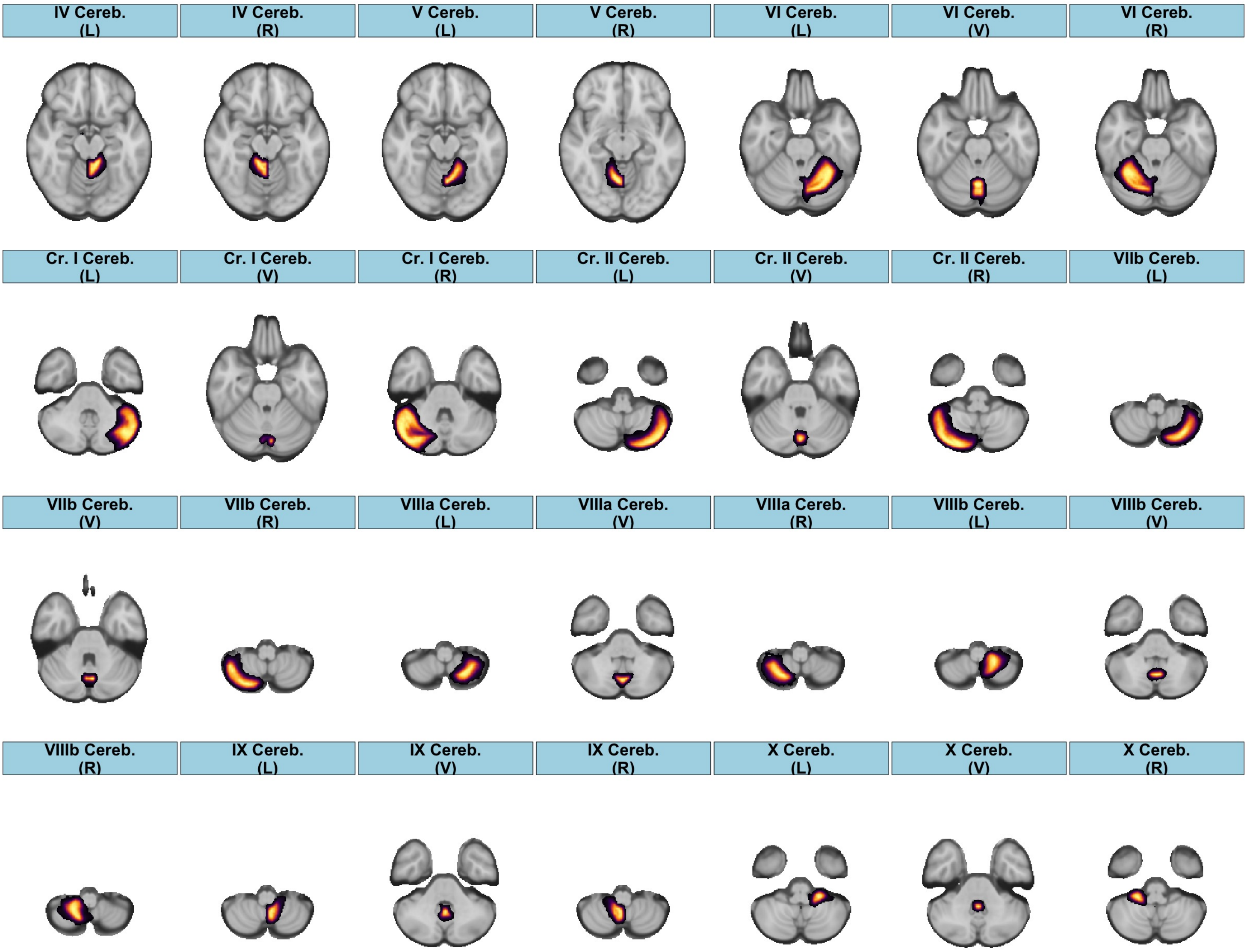


*Figure S3. The 28 ROIs from the Diedrichsen structural atlas.*

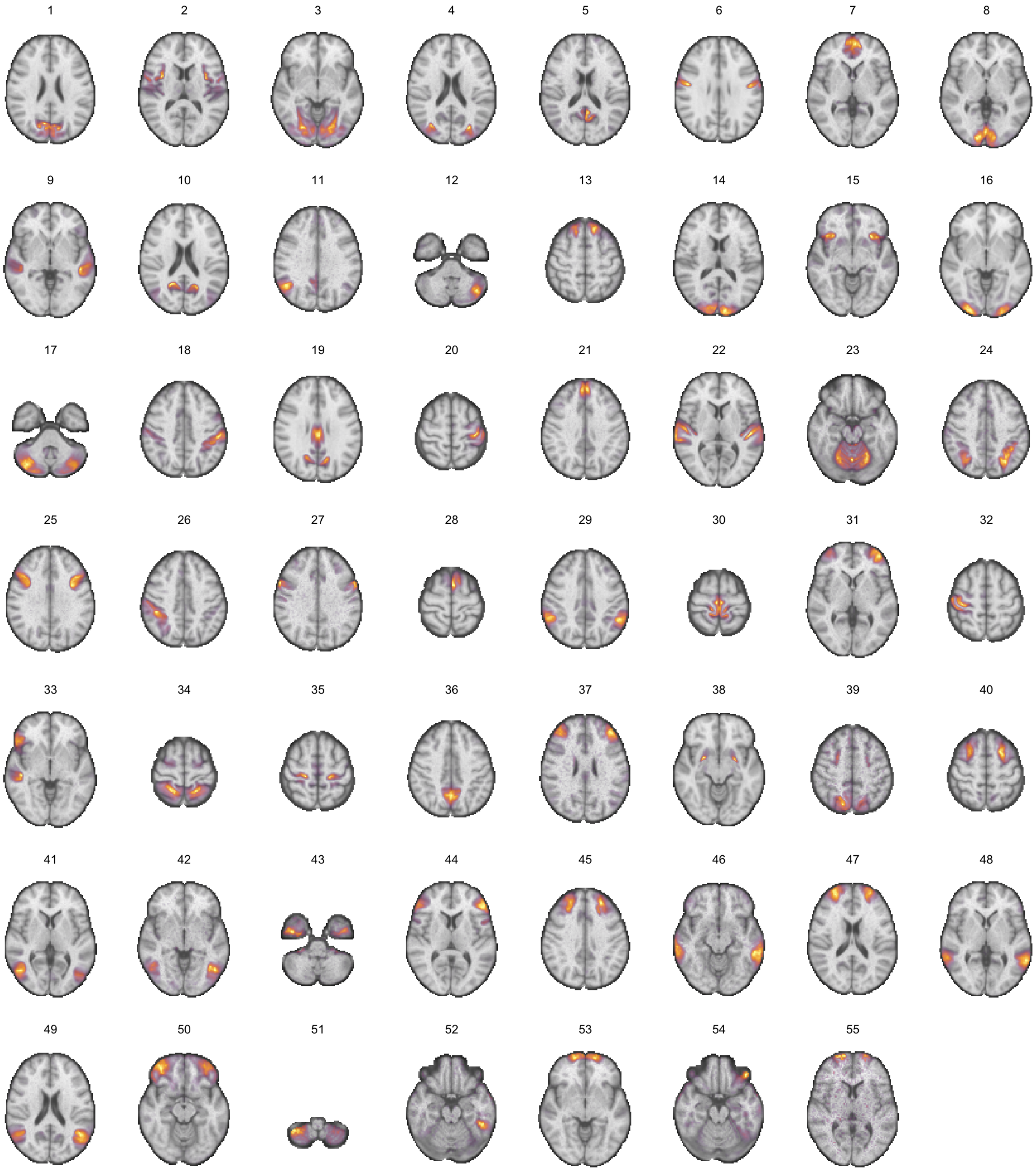


*Figure S4. The 55 non-artefact components from parcellations of the brain obtained from the rs-fMRI pipeline in UKB.*

### 4. Flow Charts for Data Processing

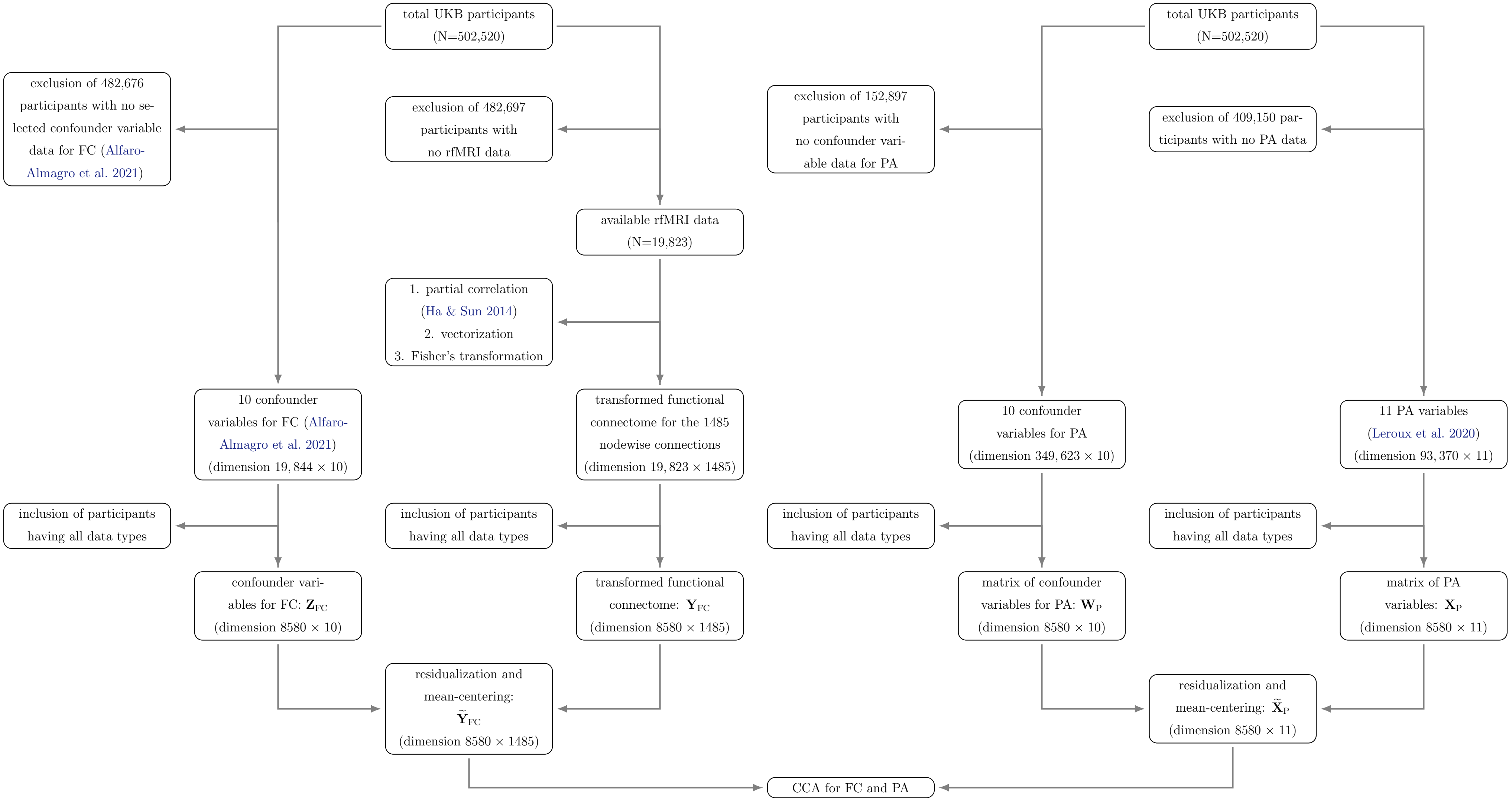


*Figure S5. Data processing flow chart of CCA relating FC to PA variables.*

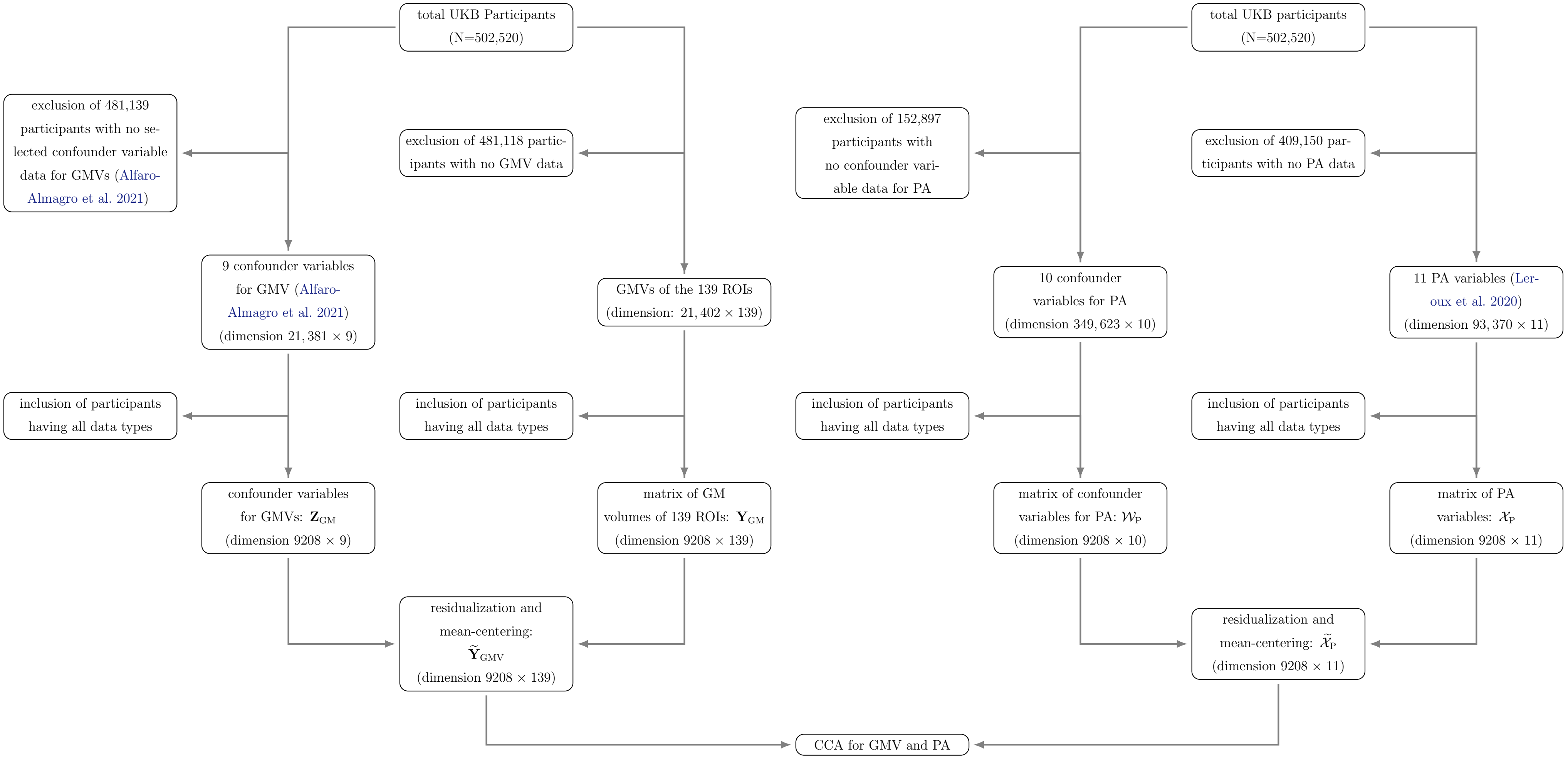


*Figure S6. Data processing flow chart of CCA relating GMV to PA variables.*

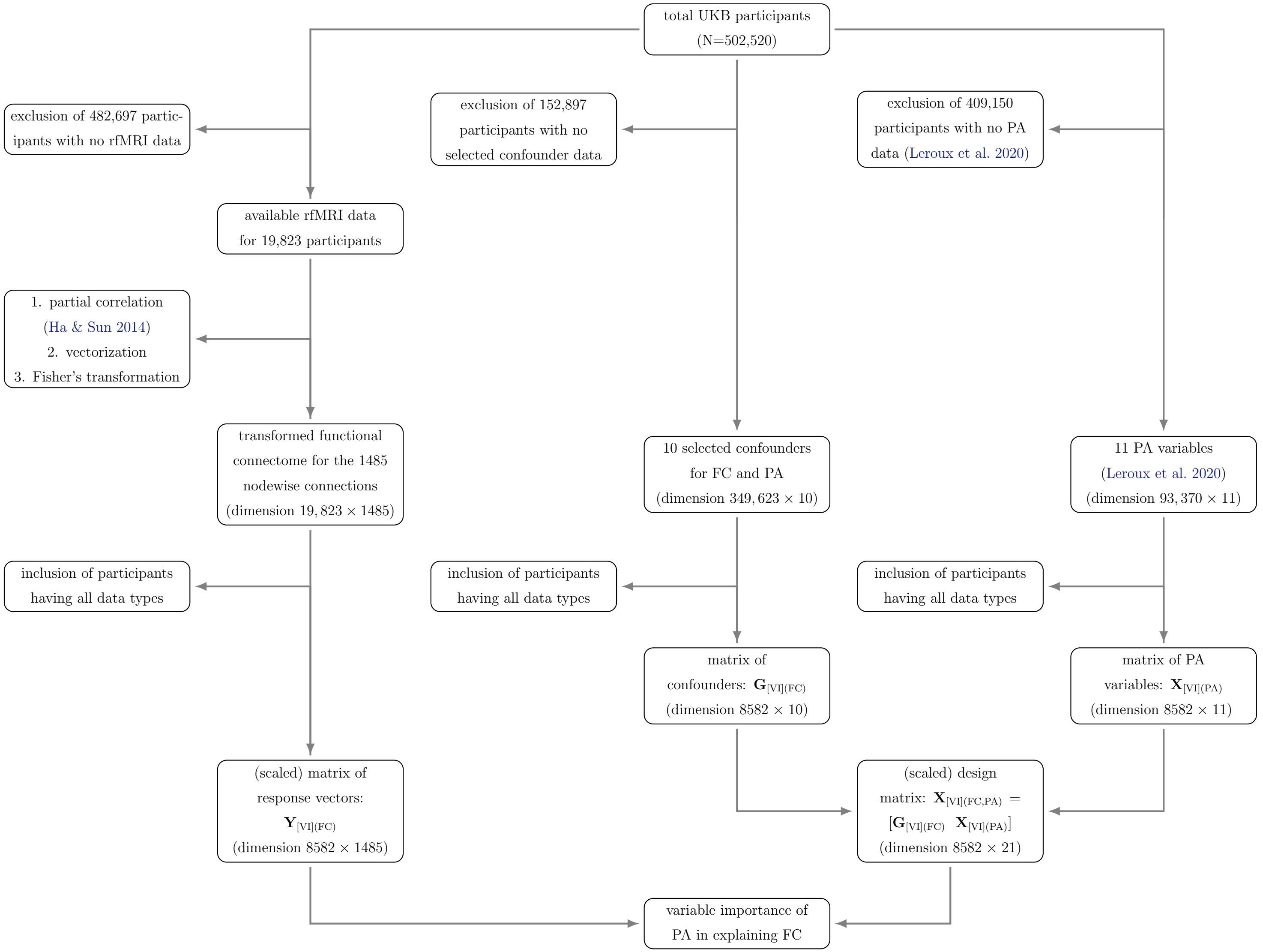


*Figure S7. Data processing flow chart for assessing variable importance (VI) of individual PA variable in modeling FC.*

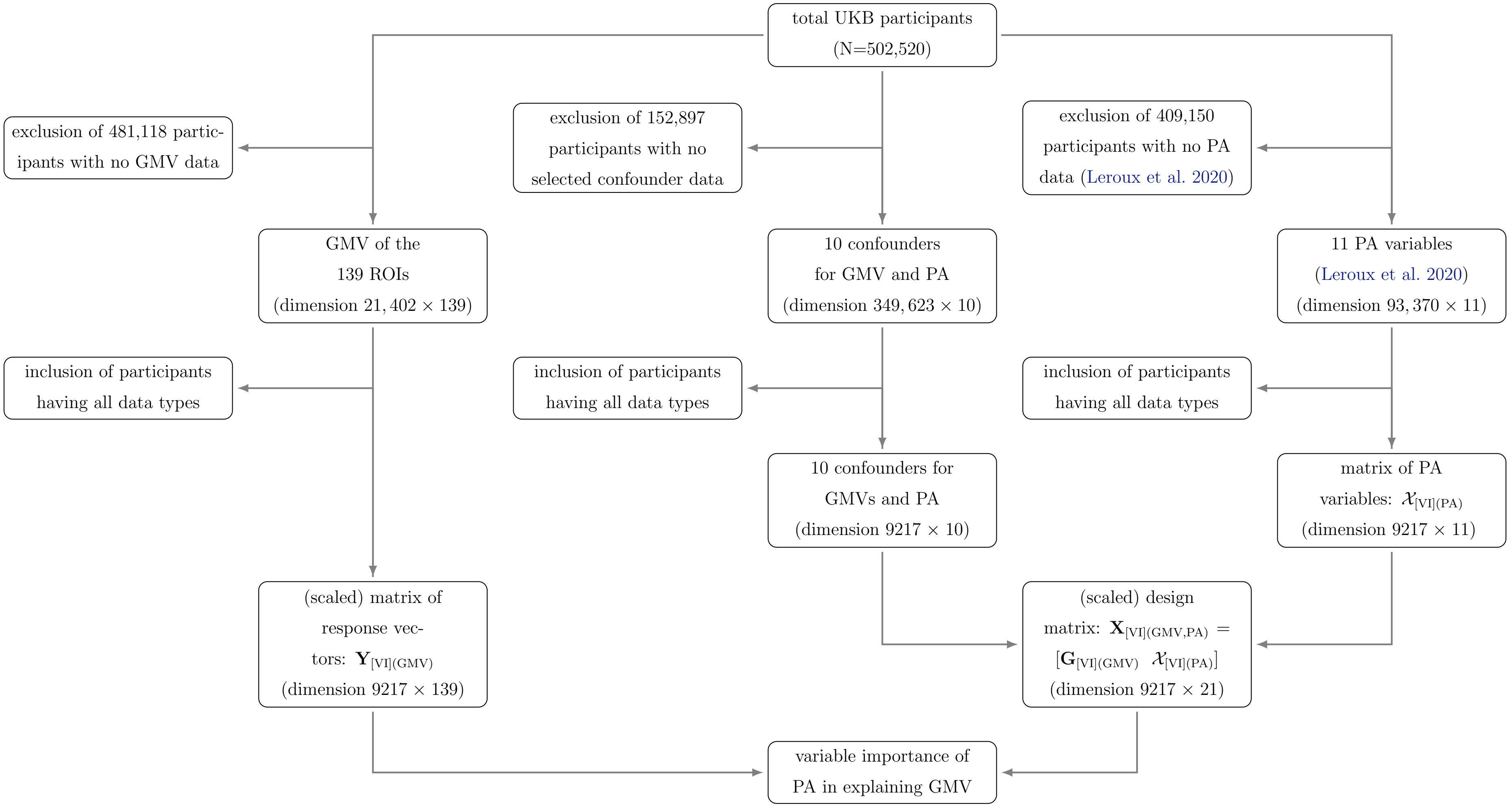


*Figure S8. Data processing flow chart for assessing variable importance (VI) of individual PA variable in modeling GMV.*

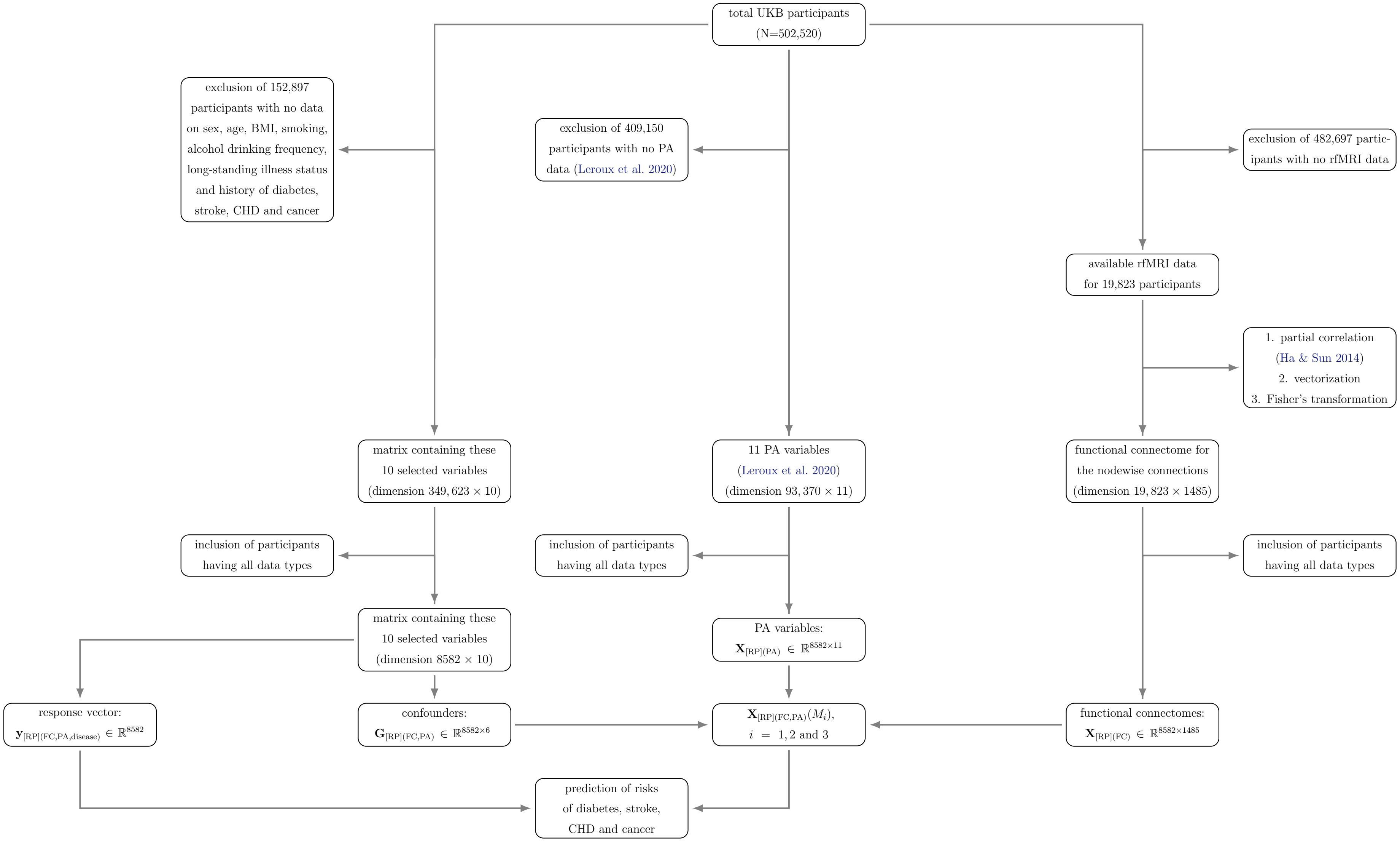


*Figure S9. Data processing flow chart for predicting the risks of diabetes, stroke, CHD and cancer via nested logistic models containing FC and PA.*

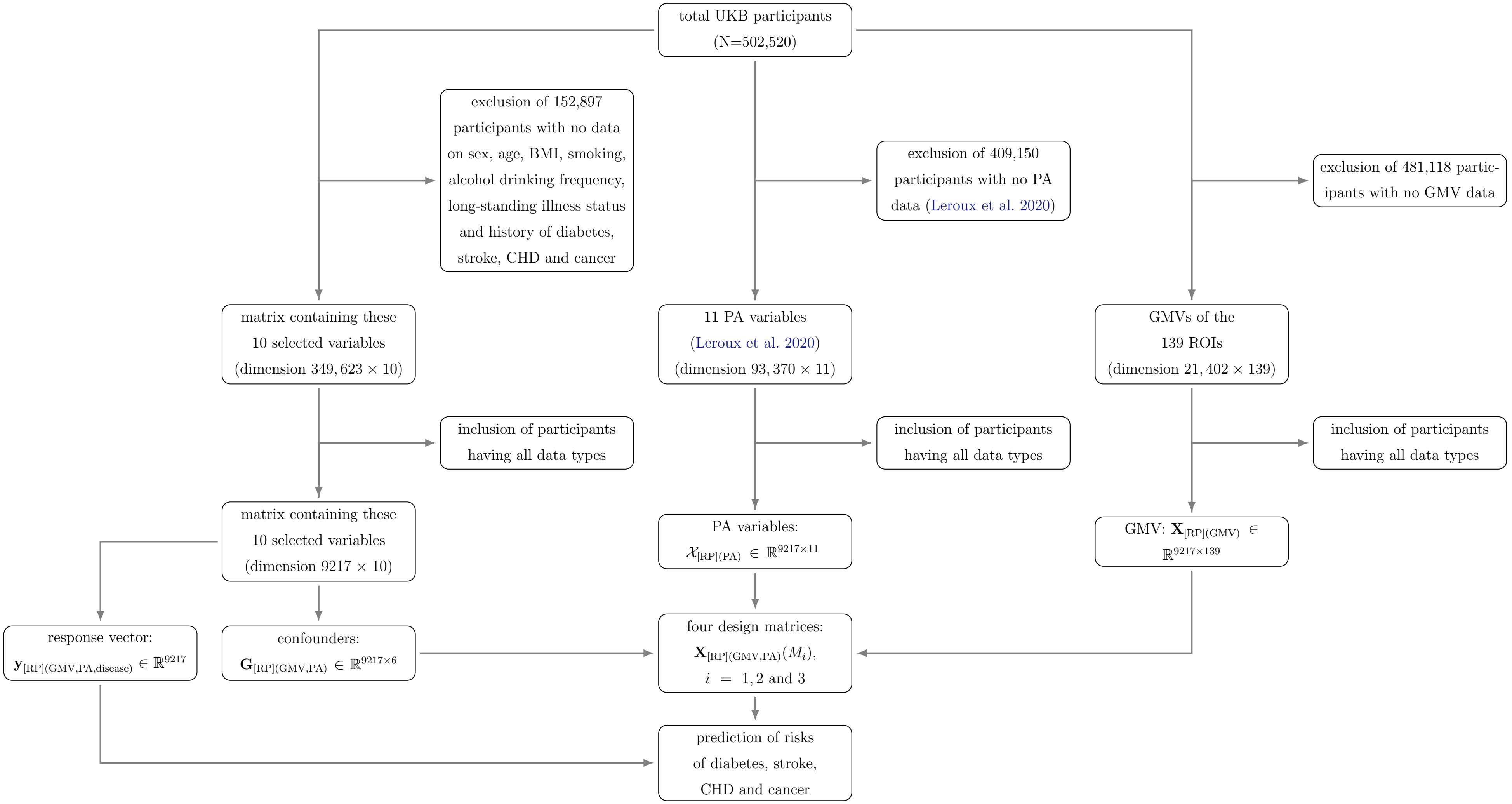


*Figure S10. Data processing flow chart for predicting the risks of diabetes, stroke, CHD and cancer by nested logistic models containing GMV and PA.*

### 5. Population-level Pattern of the Functional Connectivity for CCA and Variable Importance Assessment

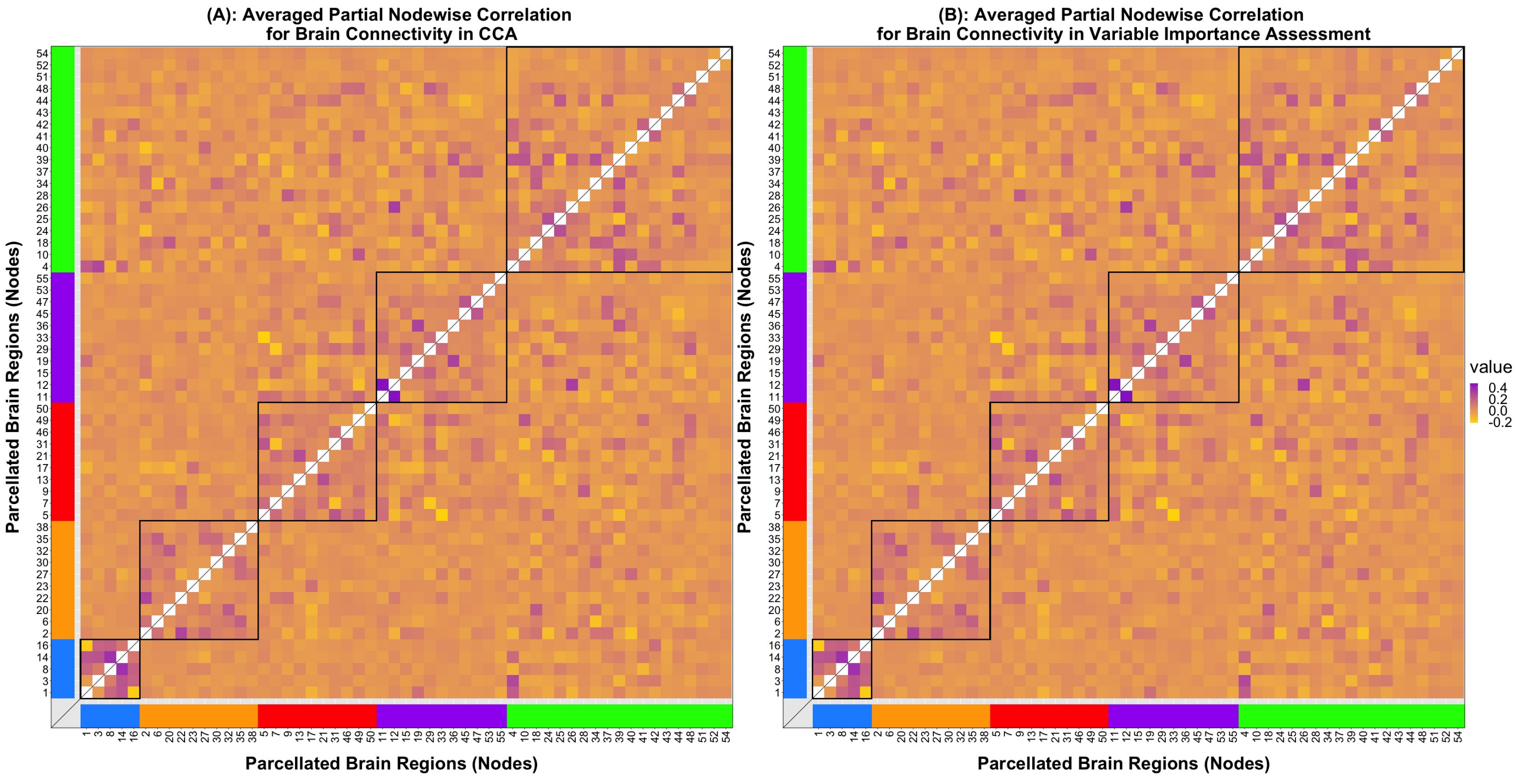


*Figure S11. (A) mean functional connectivity matrix used in CCA relating FC to PA variables; (B) mean functional connectivity matrix used in assessing the importance of individual PA variable when modeling FC. The parcellated brain regions (nodes) are grouped by the five resting-state networks (RSNs). Specifically, the blue, orange, red, purple and green networks respectively controls visual, sensorimotor, default, cingulo-opercular, and control and attention functions. The diagonal elements are omitted.*

### 6. Testing of Significance of Linear Regression Models in Estimating the GMVs of the 139 ROIs

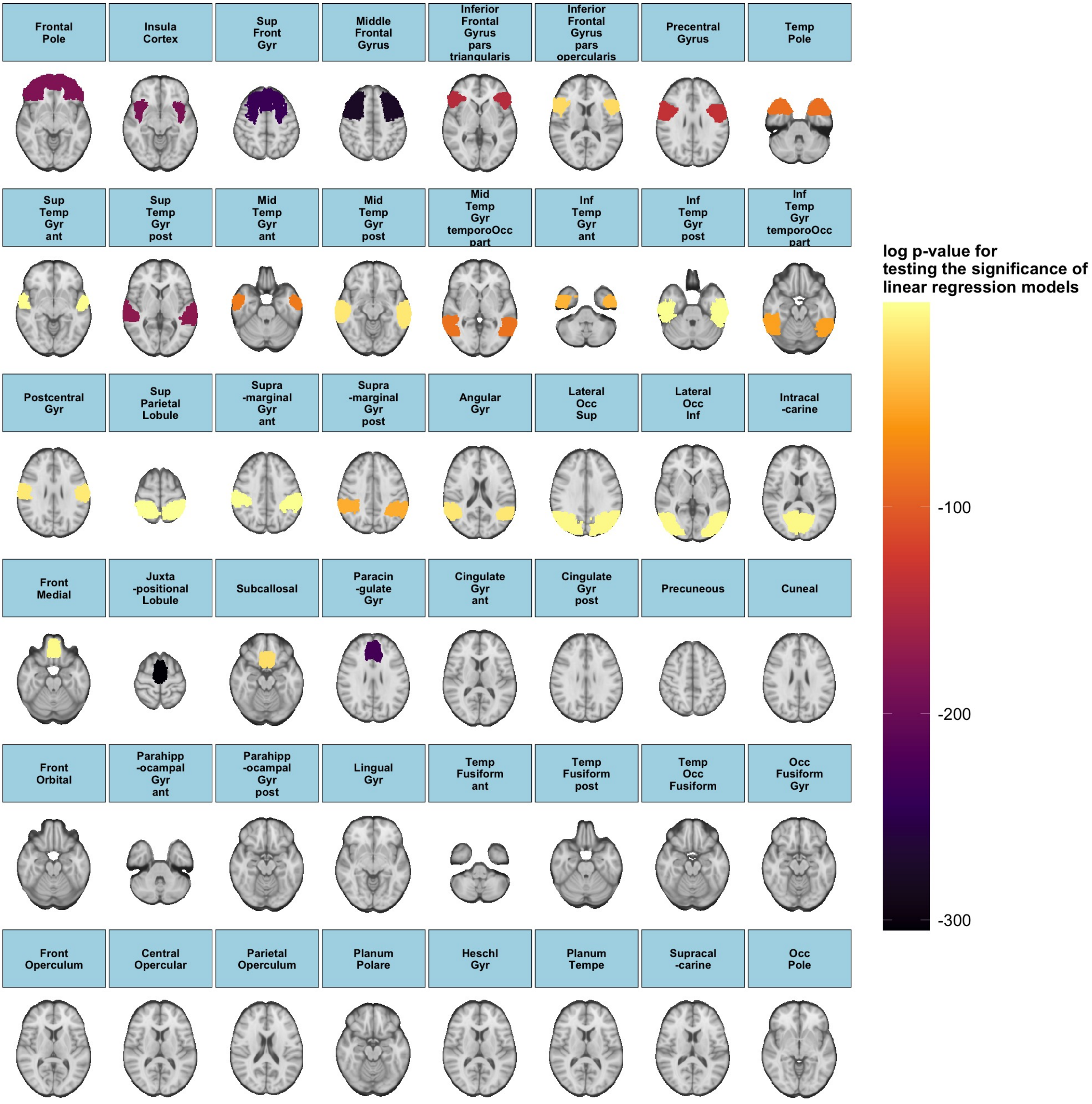


*Figure S12. The logarithm of adjusted p-values associated with testing the significance of linear regression models in estimating the GMVs of the 96 ROIs from the Harvard-Oxford cortical structural atlas.*

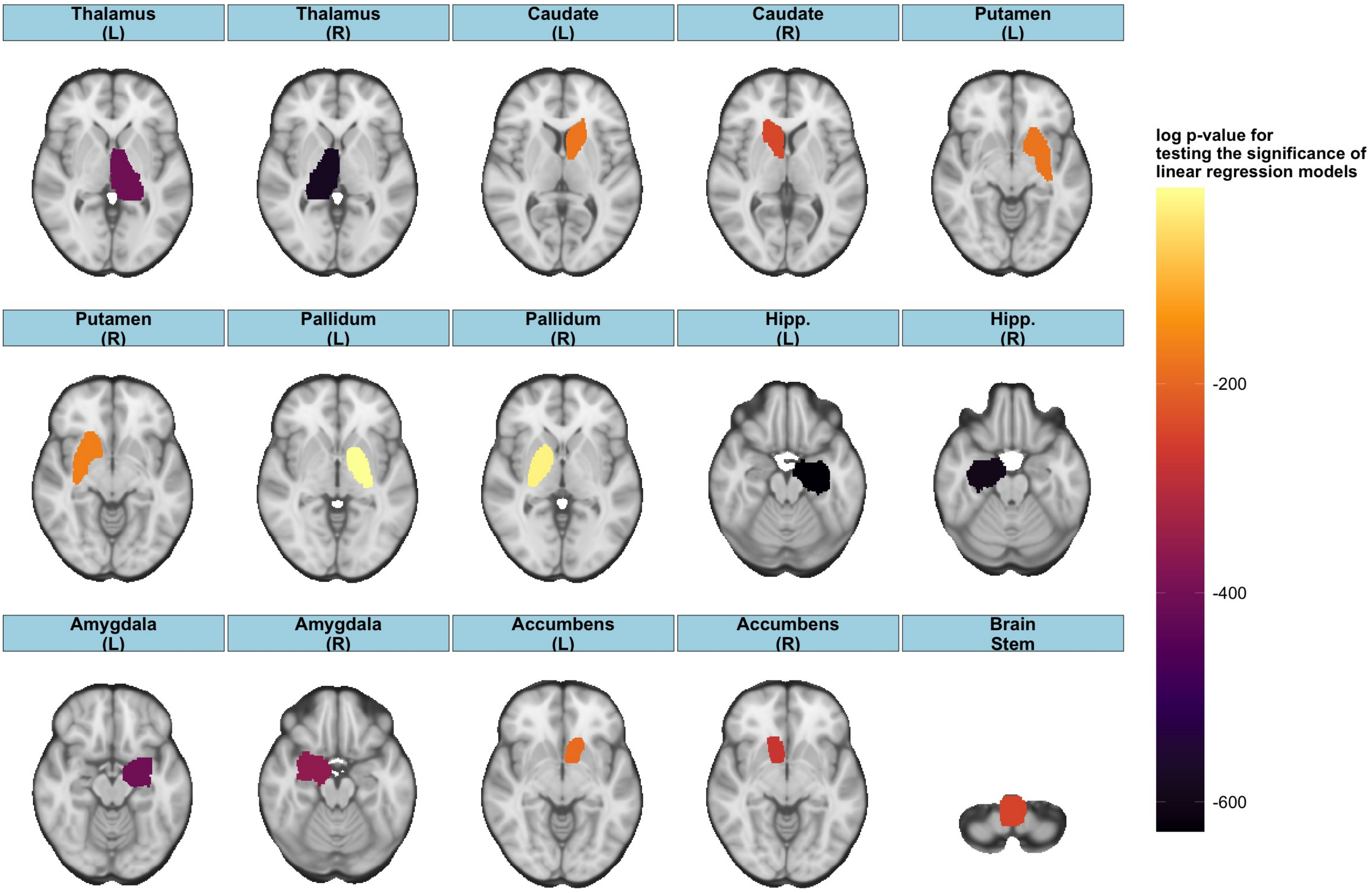


*Figure S13. The logarithm of adjusted p-values associated with testing the significance of linear regression models in estimating the GMVs of the 15 ROIs from the Harvard-Oxford subcortical structural atlas.*

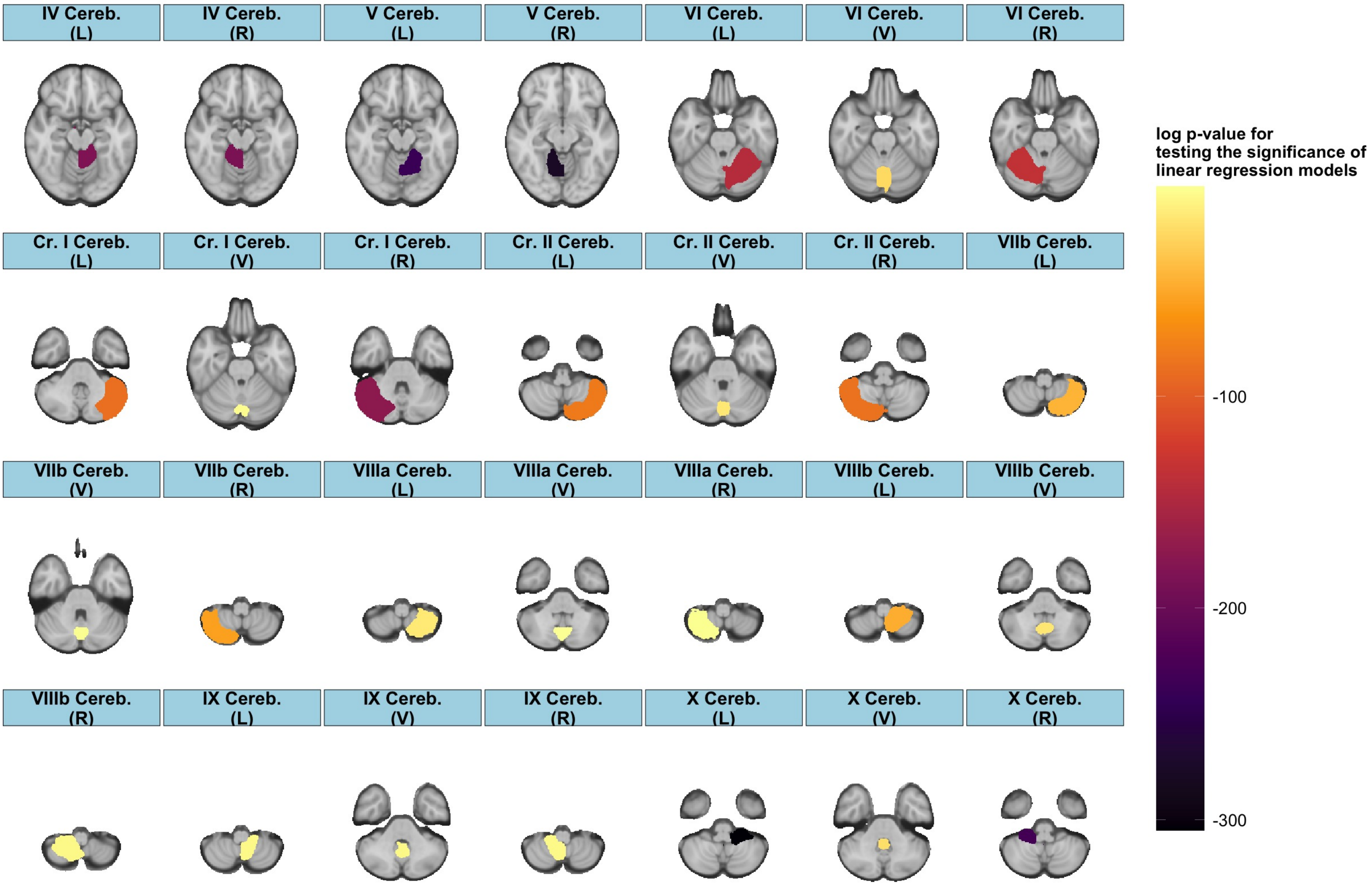


*Figure S14. The logarithm of adjusted p-values associated with testing the significance of linear regression models in estimating the GMVs of the 28 ROIs from the Diedrichsen structural atlas.*

### 7. Method for Canonical Correlation Analysis (CCA)

**Population CCA**. CCA (Hotelling 1936) is a multivariate statistical inference method that quantifies the association between two random vectors $\boldsymbol{x} \in \mathbb{R}^p$ and $\boldsymbol{y} \in \mathbb{R}^q$, where $p$ does not necessarily equal $q$. The approach seeks to identify $k$ sets of vectors $\boldsymbol{a}_i \in \mathbb{R}^p$ and $\boldsymbol{b}_i \in \mathbb{R}^q$, $i = 1, ..., k$ where $k = \min(p, q)$, that sequentially maximize the Pearson's correlation between the linear combinations $\boldsymbol{a}_i^\top \boldsymbol{x}$ and $\boldsymbol{b}_i^\top \boldsymbol{y}$. Mathematically, such sequence of vectors $\boldsymbol{a}_i$ and $\boldsymbol{b}_i$ are obtained by solving:

$$(\boldsymbol{a}_i, \boldsymbol{b}_i) = \underset{\boldsymbol{\alpha} \in \mathbb{R}^p, \boldsymbol{\beta} \in \mathbb{R}^q}{\operatorname{argmax}} \operatorname{Corr}(\boldsymbol{\alpha}^\top \boldsymbol{x}, \boldsymbol{\beta}^\top \boldsymbol{y}),\ i = 1, ..., k, \tag{1}$$

subject to the constraints that for $i, j = 1, ..., k$,

$$(1)\ \operatorname{Cov}(\boldsymbol{a}_i^\top \boldsymbol{x}, \boldsymbol{a}_j^\top \boldsymbol{x}) = \operatorname{Cov}(\boldsymbol{b}_i^\top \boldsymbol{y}, \boldsymbol{b}_j^\top \boldsymbol{y}) = \operatorname{Cov}(\boldsymbol{a}_i^\top \boldsymbol{x}, \boldsymbol{b}_j^\top \boldsymbol{y}) = 0,\ i \neq j, \text{ and} \tag{2}$$

$$(2)\ \rho_1 \geq ... \geq \rho_k \text{ with } \rho_i = \operatorname{Corr}(\boldsymbol{a}_i^\top \boldsymbol{x}, \boldsymbol{b}_i^\top \boldsymbol{y}), \tag{3}$$

where $\operatorname{Corr}(\cdot, \cdot)$ and $\operatorname{Cov}(\cdot, \cdot)$ represent the correlation and covariance between two random variables, respectively. Hence, CCA successively optimizes the correlation between linear combinations of $\boldsymbol{x}$ and $\boldsymbol{y}$ with the following requirements: $(i)$ different pairs of linear combinations of $\boldsymbol{x}$, $\boldsymbol{y}$, and $\boldsymbol{x}$ and $\boldsymbol{y}$ are uncorrelated, and $(ii)$ the sequence of vectors $\boldsymbol{a}_i$ and $\boldsymbol{b}_i$ is obtained such that the correlation of the corresponding pairs of linear combinations of $\boldsymbol{x}$ and $\boldsymbol{y}$ are in a non-increasing manner.

The terms $\boldsymbol{a}_i$ and $\boldsymbol{b}_i$ are called the $i^{\text{th}}$ canonical coefficients, and the pair $U_i = \boldsymbol{a}_i^\top \boldsymbol{x}$ and $V_i = \boldsymbol{b}_i^\top \boldsymbol{y}$ the $i^{\text{th}}$ canonical variates. Moreover, $\rho_i$ defined in (3) is called the canonical correlation. Canonical correlations are given by $\rho_i = \sqrt{\lambda_i}$, $i = 1, ..., k$, where $\lambda_1 \geq ... \geq \lambda_k$ are the $k$ largest eigenvalues of $\Sigma_1 = \Sigma_{\boldsymbol{y}}^{-1/2} \Sigma_{\boldsymbol{yx}} \Sigma_{\boldsymbol{x}}^{-1} \Sigma_{\boldsymbol{xy}} \Sigma_{\boldsymbol{y}}^{-1/2}$, which are identical to the $k$ largest eigenvalues of $\Sigma_2 = \Sigma_{\boldsymbol{x}}^{-1/2s} \Sigma_{\boldsymbol{xy}} \Sigma_{\boldsymbol{y}}^{-1} \Sigma_{\boldsymbol{yx}} \Sigma_{\boldsymbol{x}}^{-1/2}$. Here, $\Sigma_{\boldsymbol{x}}$ and $\Sigma_{\boldsymbol{y}}$ are the variances of $\boldsymbol{x}$ and $\boldsymbol{y}$, and $\Sigma_{\boldsymbol{xy}} = \Sigma_{\boldsymbol{yx}}^\top$ is the covariance matrix of $\boldsymbol{x}$ and $\boldsymbol{y}$. Let $\boldsymbol{\kappa}_i$ and $\boldsymbol{\nu}_i$ denote the eigenvectors of $\Sigma_1$ and $\Sigma_2$ corresponding to $\lambda_i$ in non-increasing order, then the canonical coefficients $\boldsymbol{a}_i$ and $\boldsymbol{b}_i$ are given by $\boldsymbol{a}_i = \boldsymbol{\nu}_i \Sigma_{\boldsymbol{x}}^{-1/2}$ and $\boldsymbol{b}_i = \boldsymbol{\kappa}_i \Sigma_{\boldsymbol{y}}^{-1/2}$, respectively.

**Sample CCA**. Given random samples $\boldsymbol{x}_i$ and $\boldsymbol{y}_i$, where $\boldsymbol{x}_i \in \mathbb{R}^p$ and $\boldsymbol{y}_i \in \mathbb{R}^q$, $i = 1, ..., n$, the sample version of CCA determines $k$ sets of vectors $\widehat{\boldsymbol{a}}_j$ and $\widehat{\boldsymbol{b}}_j$ for $j = 1, ..., k$ where $k = \min(p, q)$ such that

$$(\widehat{\boldsymbol{a}}_j, \widehat{\boldsymbol{b}}_j) = \underset{\widetilde{\boldsymbol{\alpha}} \in \mathbb{R}^p, \widetilde{\boldsymbol{\beta}} \in \mathbb{R}^q}{\operatorname{argmax}} \widehat{\operatorname{Corr}}(\mathbf{X}\widetilde{\boldsymbol{\alpha}}, \mathbf{Y}\widetilde{\boldsymbol{\beta}}),\ j = 1, ..., k,$$

subject to the constraints that for $l, m = 1, ..., k$,

$$(1)\ \widehat{\text{Cov}}(\mathbf{X}\widehat{\boldsymbol{a}}_l, \mathbf{X}\widehat{\boldsymbol{a}}_m) = \widehat{\text{Cov}}(\mathbf{Y}\widehat{\boldsymbol{b}}_l, \mathbf{Y}\widehat{\boldsymbol{b}}_m) = \widehat{\text{Cov}}(\mathbf{X}\widehat{\boldsymbol{a}}_l, \mathbf{Y}\widehat{\boldsymbol{b}}_m) = 0,\ l \neq m,\ \text{and}$$

$$(2)\ \widehat{\rho}_1 \geq ... \geq \widehat{\rho}_k,\ \rho_l = \widehat{\text{Corr}}(\mathbf{X}\widehat{\boldsymbol{a}}_l, \mathbf{Y}\widehat{\boldsymbol{b}}_l),$$

where $\widehat{\text{Corr}}(\cdot, \cdot)$ and $\widehat{\text{Cov}}(\cdot, \cdot)$ are the sample correlation and covariance between two sample vectors, and $\mathbf{X} = (\boldsymbol{x}_1, ..., \boldsymbol{x}_n)^\top \in \mathbb{R}^{n\times p}$ and $\mathbf{Y} = (\boldsymbol{y}_1, ..., \boldsymbol{y}_n)^\top \in \mathbb{R}^{n\times q}$ refer to the $n \times p$ and $n \times q$ matrices resulting from row-wise concatenation of $\boldsymbol{x}_i$'s and $\boldsymbol{y}_i$'s, respectively. The sample canonical coefficients are given by $\widehat{\rho}_j = \sqrt{\widehat{\lambda}_j}$, $j = 1, ..., k$, where $\widehat{\lambda}_1 \geq ... \geq \widehat{\lambda}_k$ are the $k$ largest eigenvalues of $\widehat{\Sigma}_1 = S_{\mathbf{Y}}^{-1/2} S_{\mathbf{YX}} S_{\mathbf{X}}^{-1} S_{\mathbf{XY}} S_{\mathbf{Y}}^{-1/2}$. These $k$ eigenvalues are the same to their counterparts of $\widehat{\Sigma}_2 = S_{\mathbf{X}}^{-1/2} S_{\mathbf{XY}} S_{\mathbf{Y}}^{-1} S_{\mathbf{YX}} S_{\mathbf{X}}^{-1/2}$, where $S_{\mathbf{X}}$ and $S_{\mathbf{Y}}$ are the sample variances of $\mathbf{X}$ and $\mathbf{Y}$, and $S_{\mathbf{XY}} = S_{\mathbf{YX}}^\top$ is the sample covariance of $\mathbf{X}$ and $\mathbf{Y}$. Moreover, the sample canonical coefficients are given by $\widehat{\boldsymbol{a}}_j = S_{\mathbf{Y}}^{-1/2}\widehat{\nu}_j$ and $\widehat{\boldsymbol{b}}_j = S_{\mathbf{X}}^{-1/2}\widehat{\kappa}_j$, where $\widehat{\nu}_j$ and $\widehat{\kappa}_j$ are the eigenvectors of $\widehat{\Sigma}_1$ and $\widehat{\Sigma}_2$ corresponding to the eigenvalues $\widehat{\lambda}_j$. As such, the $j^{\text{th}}$ sample canonical variates are $\widehat{\mathbf{U}}_j = \mathbf{X}S_{\mathbf{Y}}^{-1/2}\widehat{\nu}_j$ and $\widehat{\mathbf{V}}_j = \mathbf{Y}S_{\mathbf{X}}^{-1/2}\widehat{\kappa}_j$.

**Bipartial CCA**. The presence of confounder variables can distort the relationship between $\mathbf{X}$ and $\mathbf{Y}$. For these reasons, CCA is often performed on data after removing the effects of confounders. In other words, it is performed on the residuals $\widetilde{\mathbf{X}}$ and $\widetilde{\mathbf{Y}}$ obtained after respectively regressing $\mathbf{W}$ and $\mathbf{Z}$ from $\mathbf{X}$ and $\mathbf{Y}$, where $\mathbf{W}$ and $\mathbf{Z}$ denote the confounder variables for $\mathbf{X}$ and $\mathbf{Y}$, respectively. The sample CCA is then used to measure the association between the residuals $\widetilde{\mathbf{X}}$ and $\widetilde{\mathbf{Y}}$, an approach known as bipartial CCA (Lee 1978). This leads to the residualized canonical correlations $\widetilde{\rho}_j$, canonical coefficients $\widetilde{\boldsymbol{a}}_j$ and $\widetilde{\boldsymbol{b}}_j$, and canonical variates $\widetilde{\mathbf{U}}_j = \widetilde{\mathbf{X}}\widetilde{\boldsymbol{a}}_j$ and $\widetilde{\mathbf{V}}_j = \widetilde{\mathbf{Y}}\widetilde{\boldsymbol{b}}_j$ obtained as discussed above for the sample CCA, where $j = 1, ..., k$ and $k = \min(p, q)$.

**Significance Tests for Bipartial CCA**. After performing CCA, we sequentially test for the significance of the CCA modes of co-variation, a term referring to the pairs of the canonical variates. This allows us to determine whether the identified canonical correlations are statistically significant. In case of bipartial CCA, this is accomplished by successively testing the null hypotheses $\widetilde{H}_0^j : \widetilde{\rho}_1 \neq 0, ..., \widetilde{\rho}_j \neq 0,\ \widetilde{\rho}_{j+1} = \cdots = \widetilde{\rho}_k = 0$, $j = 1, ..., k$. A possible test statistic is $\widetilde{\zeta}_j = -(n - 1 - (p + q + 1)/2) \log \left(\prod_{i=j}^k (1 - \widetilde{\rho}_i^2)\right)$ (Wilks 1935), which can be interpreted as a truncated version of the likelihood ratio test statistic. Under $\widetilde{H}_0^j$, $\widetilde{\zeta}_j$ approximately follows a $\chi^2$ distribution with $(p-j)(q-j)$ degrees of freedom, a result that hinges on large sample size $n$ and the assumption that columns of $\mathbf{X}$ and $\mathbf{Y}$ are *i.i.d.* with normal distribution. The widely-adopted permutation test, which depends on the exchangeability of the observations, serves as a plausible alternative. Specifically, randomly permuting the rows of $\mathbf{X}$ and $\mathbf{Y}$ $B$ times generates $B$ copies of the test statistic for each $\widetilde{\zeta}_j$, denoted $\widetilde{\zeta}^*_{j,1}, ..., \widetilde{\zeta}^*_{j,B}$. The $p$-value for testing $\widetilde{H}^j_0$ is computed as $\widetilde{p}_j = (1/B) \sum_{b=1}^{B} \mathbb{1}(\widetilde{\zeta}_j \geq \widetilde{\zeta}^*_{j,b})$, where $\mathbb{1}(\cdot)$ is the indicator function. This is followed by control of the false discovery rate for all of the $\widetilde{p}_j$'s (Benjamini & Hochberg 1995).

In the presence of confounder variables where bipartial CCA is utilized, it has been argued that the permutation test is susceptible to the violation of the exchangeability assumption due to the aforementioned residualization, potentially resulting in hypothesis tests which do not achieve nominal type I/II error rates for testing the significance of all the CCA modes except for the first (primary) mode (Winkler et al. 2020). To address this issue, $\mathbf{X}$ and $\mathbf{Y}$ are projected to lower-dimensional space where exchangeability holds for the permutation test. Such projection matrices for $\mathbf{X}$ and $\mathbf{Y}$ are respectively given by $\mathbf{Q_W P_X Q_W^\top}$ and $\mathbf{Q_Z P_Y Q_Z^\top}$, where $\mathbf{Q_W}$ and $\mathbf{Q_Z}$ are the semi-orthogonal bases for the column spaces of $\mathbf{I} - \mathbf{W}(\mathbf{W}^\top\mathbf{W})^{-1}\mathbf{W}^\top$ and $\mathbf{I} - \mathbf{Z}(\mathbf{Z}^\top\mathbf{Z})^{-1}\mathbf{Z}^\top$, and $\mathbf{P_X}$ and $\mathbf{P_Y}$ are known as the selection matrices, which are modified versions of the identity matrices; guidelines for their construction is available in Winkler et al. (2014). While this approach has desirable theoretical properties, numerical challenges in the computation of $\mathbf{Q_W}$ and $\mathbf{Q_Z}$ were encountered in our particular study, specifically singularity issues of $\mathbf{W}$ and $\mathbf{Z}$. Thus, in our study, we focus on analyzing the primary CCA mode of co-variation.

**Additional Statistical Analyses.**In addition to estimating canonical correlations and coefficients and performing the test of significance of the CCA modes, we report two additional results from CCA. First, we computed the cumulative proportions of the sample variance in $\widetilde{\mathbf{X}}$ and $\widetilde{\mathbf{Y}}$ explained by the canonical variates $\widetilde{\mathbf{U}}_j$ and $\widetilde{\mathbf{V}}_j$, respectively, where the individual proportion is given by $\sum_{j=1}^{k} \sum_{m=1}^{p} \widehat{\text{Corr}}(\widetilde{\mathbf{U}}_j, \widetilde{\mathbb{X}}_m)^2/p$ and $\sum_{j=1}^{k} \sum_{l=1}^{q} \widehat{\text{Corr}}(\widetilde{\mathbf{V}}_j, \widetilde{\mathbb{Y}}_l)^2/q$. Here, $\widetilde{\mathbb{X}}_m$ and $\widetilde{\mathbb{Y}}_l$ denote the $m^{\text{th}}$ and $l^{\text{th}}$ columns of $\widetilde{\mathbf{X}}$ and $\widetilde{\mathbf{Y}}$, $m = 1, ..., p$ and $l = 1, ...., q$. These quantities can be interpreted as the relative contribution of individual CCA modes in estimating (co)variability in the data. Second, we compute the sample correlation between each column of $\widetilde{\mathbf{X}}$ and the canonical variate for $\widetilde{\mathbf{Y}}$ corresponding to the primary CCA mode. This metric is calculated to assess the direction and strength of association between each PA variable and the primary CCA mode associated with FC or GMV.
